# Clonal lineage tracing of innate immune cells in human cancer

**DOI:** 10.1101/2025.07.16.665245

**Authors:** Vincent Liu, Katalin Sandor, Patrick K. Yan, Max Miao, Yajie Yin, Robert R. Stickels, Andy Y. Chen, Kamir Hiam-Galvez, Jacob Gutierrez, Wenxi Zhang, Sairaj M. Sajjath, Raeline Valbuena, Steven Wang, Bence Dániel, Leif S. Ludwig, Brooke E. Howitt, Caleb Lareau, Ansuman T. Satpathy

**Affiliations:** Department of Genetics, Stanford University, Stanford, CA, USA; Department of Pathology, Stanford University, Stanford, CA, USA; Center for Immunotherapy Design, Stanford University, Stanford, CA, USA; Department of Bioengineering, Stanford University, Stanford, CA, USA; Computational and Systems Biology Program, Memorial Sloan Kettering Cancer Center, New York, NY, USA; Robin Chemers Neustein Laboratory of Mammalian Cell Biology and Development, Howard Hughes Medical Institute, The Rockefeller University, New York, NY 10065, USA; Parker Institute for Cancer Immunotherapy, San Francisco, CA, USA; Berlin Institute of Health at Charité – Universitätsmedizin Berlin, Berlin, Germany; Max-Delbrück-Center for Molecular Medicine in the Helmholtz Association (MDC) Berlin Institute for Medical Systems Biology (BIMSB), Berlin, Germany

## Abstract

Innate immune cells constitute the majority of the tumor microenvironment (TME), where they mediate both natural anti-tumor immunity and immunotherapy responses. While single-cell T- and B-cell receptor sequencing has provided fundamental insights into the clonal dynamics of human adaptive immunity, the lack of appropriate tools has precluded similar analysis of innate immune cells. Here, we describe a method that leverages somatic mitochondrial DNA (mtDNA) mutations to reconstruct clonal lineage relationships between single cells across cell types in native human tissues. We jointly sequenced single-cell transposase-accessible chromatin and mtDNA to profile *n*=124,958 cells from matched tumor, non-involved lung tissue (NILT), and peripheral blood of early-stage non-small cell lung cancer (NSCLC) patients, as well as *n*=93,757 cells from matched tumor and peripheral blood of ovarian cancer patients. Single-cell concomitant profiling of lineage and cell states of thousands of immune cells resolved clonality across cell types, tissue sites, and malignancies. Clonal tracing of innate immune cells demonstrates that TME-resident myeloid subsets, including macrophages and type 3 dendritic cells (DC3), are clonally linked to both circulating and tissue-infiltrating monocytes. Further, we identify distinct DC-biased and macrophage-biased myeloid clones, enriched in the tumor and NILT, respectively, and find that their circulating monocyte precursors exhibit distinct epigenetic profiles, suggesting that myeloid differentiation fate may be predetermined before TME infiltration. These results delineate the clonal pathways of intratumoral myeloid cell recruitment and differentiation in human cancer and suggest that remodeling of the tumor myeloid compartment may be peripherally programmed.

## Introduction

The TME is a dynamic ecosystem that maintains a close connection to the peripheral immune system through the bloodstream. Circulating immune cells continually infiltrate the TME, where their capacity to differentiate and expand shapes anti-tumor immunity and therapy response. Lineage tracing methods such as T cell paired single-cell RNA and TCR sequencing (scRNA/TCR-seq) have advanced understanding of the expansion potential, tissue distribution, and clonal dynamics of immune populations in patients^1,2^. For example, we and others have applied scRNA/TCR-seq to show that peripherally expanded, tumor-specific CD8^+^ T cell clones replace dysfunctional exhausted T cells in the tumor, which is essential to immune checkpoint blockade response^3–5^. However, similar lineage-tracing approaches have not been feasible for non-adaptive immune populations due to the absence of somatic recombination events during their development.

Innate immune cells, including myeloid cells, are among the most abundant immune cell types in the TME^6^. These cells are ontogenically diverse, consisting of both tissue-resident and circulating bone marrow-derived tissue-infiltrating compartments^7^. Detailed phenotypic and quantitative maps of innate immune populations across tissue sites and cancer types have delineated their phenotypic diversity,^8–11^ and there is growing recognition that ontogeny plays a key role in shaping their function within the TME^12,13^. However, due to the inability to connect cellular phenotype and lineage within native human tissues, several open questions remain. First, do innate immune cells have an intrinsic tissue site preference for infiltration, differentiation, or expansion? Second, do innate immune cells face clonally selective pressures in the TME, analogous to those experienced by T and B cells, or does their presence in tumors depend on broader cellular features independent of clonality? Finally, what specific lineage relationships connect the diverse innate immune constituents within the TME, and what are their ontogenetic origins?

Somatic mtDNA variants have been established as an endogenous lineage marker with the high mutation rate of mtDNA compared to nuclear DNA, favoring the accumulation of distinguishing mutations within cell lineages over time^14,15^. Recent advancements in single-cell sequencing have enabled the detection of mtDNA mutations^16–19^, enabling unbiased, simultaneous lineage and epigenetic profiling of thousands of clonotypes across cell types, tissue sites, and cancer types. Therefore, to address these fundamental questions regarding innate immune cell clonality and ontogeny, we performed mitochondrial single-cell Assay for Transposase Accessible Chromatin by sequencing (mtscATAC-seq) on 218,715 cells from tumor, non-involved lung tissue (NILT), and/or peripheral blood mononuclear cells (PBMCs) from patients with lung or ovarian cancer. We developed a computational framework, termed Mitotrek, to bolster the accuracy of lineage tracing analyses from mtDNA by prioritizing the fidelity of clonal assignments over detection sensitivity. By mapping the lineage relationships of myeloid cells across tissue sites, we found no evidence of clonal selection or tissue infiltration bias. However, we identified distinct clones biased towards dendritic cell (DC) or macrophage cell states, along with associated epigenetic signatures that differentiate monocytes within these groups. Overall, our work provides a clonally resolved view of the human innate immune response to cancer, with the potential to inform the development of myeloid-targeted immunotherapies.

### Development of Mitotrek to recover single-cell clonal fractions

To establish a lineage tracing method applicable to donor-matched solid tissue and blood samples and to clonally link immune cells across tissue sites, we applied mtscATAC-seq to profile five patients with early-stage non-small cell lung cancer (NSCLC) who had undergone therapeutic lobectomy (**Figure 1A**; **Table S1** and **Methods**). Three patients diagnosed with adenocarcinoma (SU-L-002, SU-L-004, and SU-L-005) were treatment-naïve, whereas one patient with squamous cell carcinoma (SU-L-001) had undergone neoadjuvant chemotherapy. A fifth patient (SU-L-003) had a neuroendocrine tumor and was receiving olaparib (PARP inhibitor) at the time of surgery for concurrent tubo-ovarian carcinosarcoma. For each patient, we collected tumor resections, non-adjacent NILT (not involved by tumor on pathological analysis), and peripheral blood samples. Fresh samples were dissociated, and single-cell ATAC and mtDNA genome-sequencing libraries were generated, enabling simultaneous recovery of chromatin accessibility profiles and mtDNA genotypes from individual cells.

**Figure 1:**
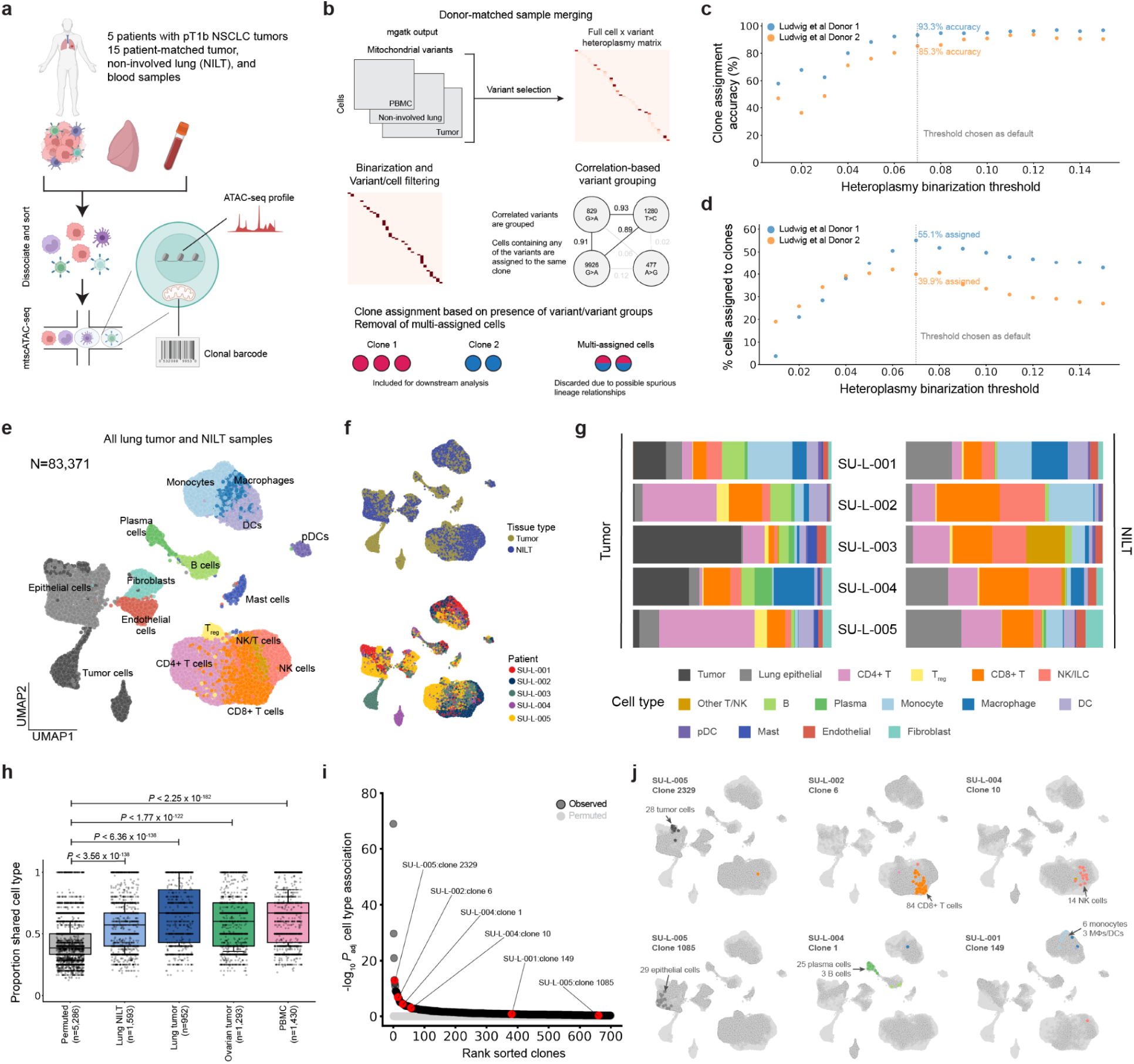
Creating a lineage-embedded atlas of NSCLC from mitochondrial DNA with Mitotrek. (A) Schematic of simultaneous single-cell epigenetic profiling and clone tracing in patients with early-stage lung adenocarcinoma. The chromatin accessibility profile and mitochondrial mutations are recovered from each cell. Created with BioRender.com. (B) Schematic of clone calling from mitochondrial variants. For each patient, mitochondrial variants from cells across all samples are used to assign cells to high-confidence clones via a stringent filtering procedure. To prioritize accuracy over sensitivity, a minimum heteroplasmy threshold is required for a variant to be considered present in a cell. Highly correlated variants, which could result from bona fide subclonal structure, were collapsed. Cells assigned to multiple clones are discarded. (C and D) Benchmarking Mitotrek using single-cell data from single-cell derived HSPC colonies in two independent donors. Clone assignment rates across tested values for the hyperparameter heteroplasmy binarization threshold, which considers heteroplasmy value below the threshold as not detected. Highest clone assignment rate was achieved at 0.07, which is chosen as the default value for the processing of all other samples in this study. (E) Uniform manifold approximation and projection (UMAP) of 83,371 cells in lung tumor and non-involved lung tissue (NILT). Cell types denoted by color are inferred after iterative sub-clustering of each of the myeloid, lymphoid, and epithelial compartments. (F) UMAP of cells colored by tissue type (top) and patient identity (bottom). (G) Normalized bar plot showing cell type composition for each patient, partitioned by tumor and non-involved lung. (H) Distribution of the proportion of cells within each clone (≥3 cells) that share the most common cell type for that clone compared to permuted data from the same samples. N=number of clones, Kruskal-Wallis test. (I) Clone associations with cell type. P-values represent the Benjamini-Hochberg adjusted Kruskal-Wallis test against overall cell type proportions for clones with at least 5 cells. Clones shown in a) are highlighted in red. (J) Representative clones capturing clonal expansion (tumor, CD8^+^ T, NK, epithelial) or differentiation (B/plasma, monocyte/macrophage) events in tumor and NILT. For each clone, cells from the clone’s donor are highlighted with shaded circles, and cells assigned to that clone are colored by their cell type.

In total, we obtained mtscATAC-seq profiles from 83,371 immune, malignant, and stromal cells from tumor and NILT, and 46,678 mtscATAC-seq profiles from the peripheral blood. Across all samples, we recovered 10,584 high-confidence somatic mtDNA mutations that are present at low pseudobulk frequencies (<1%) as determined by *mgatk*,^17^ a computational pipeline to process mitochondrial reads and generate heteroplasmy estimation from mtscATAC-seq data (**Figure S1A**). Importantly, somatic mtDNA variants have been demonstrated to arise in hematopoietic stem cells^16^ and to serve as particularly useful lineage markers to study immune dynamics, including in the innate immune system^20^. In contrast, mtDNA variants with >1% pseudobulk frequencies, accounting for 0.76% all variants detected by *mgatk*, are broadly distributed across cell types and tissue sites, likely representing early-developmental/zygotic mutations^21^ not informative for resolving more recent immune lineage relationships (**Figure S1B**). Notably, despite their relatively high pseudobulk frequencies, these variants may exhibit low heteroplasmy or be absent in cells that genuinely belong to the associated lineages due to random genetic drift from cell divisions^22^, limiting their utility in reconstructing clonal relationships. Thus, we excluded variants with >1% pseudobulk frequencies from downstream analysis.

Analysis of heteroplasmy distributions revealed that mtDNA variants detected at high allele frequencies in solid tissue cells were more frequently shared among cells, including from other lineages, compared to cells from the matching patient PBMC sample, likely due to both technical artifacts (e.g., uptake of ambient mtDNA during single-cell capture^23^) and biological phenomena (e.g., horizontal mitochondrial transfer events^24^) that are unlikely to reflect *bona fide* clonal relationships (**Figures S1C-S1H**). Notably, such issues have not been reported in prior mtDNA-lineage tracing studies of PBMCs or bone marrow, indicating they may be unique challenges in solid tumor profiling^18,20,25^. To mitigate such spurious lineage inferences, we developed a computational framework, Mitotrek, which assigns cells to high-confidence clones by prioritizing clonal accuracy over sensitivity (**Methods**). Briefly, Mitotrek: 1) reformats cell-by-variant heteroplasmy data into a binary “positive-unlabeled” matrix using empirically defined thresholds to account for measurement noise from genetic drift^22^ and variable sequencing depth; 2) discards variants detected in >20% of cells within a sample, which are likely to generate artifactual linkages, and 3) excludes cells assigned to multiple clones to avoid contamination by non-informative variants of technical or biological origin. We decided to not explore subclonal structure due to the difficulty establishing true clonal hierarchy and therefore collapsed potential subclonal relationships to single clones by merging highly correlated variants (**Figure 1B**; **Methods**). We benchmarked Mitotrek on full-length scRNA-seq data from single-colony hematopoietic stem and progenitor cells (HSPCs) with known clonal identities^16^, achieving assignment accuracies of 91.3% and 85.3% for two independent donors (**Figures 1C, 1D, S1I**, and **S1J**). Given the conservative premises of Mitotrek, clone assignment rates were lower (39.9% and 55.1%) compared to the original study (∼80%) which relied solely on the presence of clone-defining variants. These results establish Mitotrek as a high-accuracy framework for resolving clonal relationships at single-cell resolution in solid human tissues, with an intentional trade-off in detection sensitivity.

### Constructing multi-modal atlases of NSCLC and ovarian cancer

We next constructed a single-cell epigenetic atlas of NSCLC by iteratively clustering scATAC-seq profiles from tumor and NILT samples. We identified 53 cell clusters, including: 9 malignant clusters (based on enrichment in tumor samples), 6 lung epithelial clusters (*EPCAM*, *KRT18*), 2 stromal clusters (*VWF*, *PECAM1*, *COL1A2*, *FBLN1*), 15 T cell clusters (*CD3D*, *CD8A*, *CD4*), 4 NK cell/ILC clusters (*NCR1*), 15 myeloid clusters (*CD14*, *LYZ*, *HLA-DQA1*, *KIT*), 1 B cell cluster (*MS4A1*, *PAX5*), and 1 plasma cell cluster (*TNFRSF17*, **Figures 1E** and **S2A**). All non-malignant cell type annotations contained cells from multiple donors, consistent with interpatient heterogeneity^26^, and cell-type assignments aligned with surface markers used for enrichment (**Figures 1F** and **S2B**). Cell-type distributions in tumor and NILT samples matched previous transcriptomic and proteomic datasets, including enrichment of B and plasma cells and depletion of NK cells in the tumor relative to NILT^8,27^ (**Figures 1G**). In patient-matched PBMCs, we recovered all major immune populations, with the expected absence of tissue-resident myeloid subsets (e.g., macrophages, mast cells), and additionally identified circulating HSPCs (**Figures S2C-F**). The comprehensive coverage of broad immune cell types across tissue sites enabled unbiased downstream lineage analysis.

Applying Mitotrek to the NSCLC dataset, we recovered 5,146 clones comprising 29,010 cells (23.2% of cells passing scATAC QC filters). Of these, 3,190 clones (62%) contained ≥3 cells and 573 clones (11%) contained ≥10 cells. The number of clones recovered per patient ranged from 440 to 2,146 and was highly correlated with the number of recovered cells per patient (R=0.90, **Figure S2G**). Clone assignment rates were consistent across cell types and tissue sites, ranging from 15–26% in tumor/NILT and 20–32% in PBMC (**Figures S2H** and **S2I**). There was no significant correlation between cell type abundance and clone assignment rate (*P*>0.07 for tumor/NILT; *P*>0.92 for PBMC), suggesting that cell type-intrinsic properties play a key role in shaping clonal structure. Globally, cells within the same clone were significantly more likely to share a cell type than randomly grouped cells across all tissues (**Figure 1H**), reflecting lineage fidelity and supporting the validity of Mitotrek-assigned clones. A clone-level cell type association analysis identified 226 clones (20% of clones with ≥5 cells) with cell-type compositions that significantly deviate from the background distribution (P_adj_<0.05, **Figure 1I**), capturing key expansion and differentiation events across immune, stromal, and epithelial compartments (**Figure 1J**). Together, these results demonstrate the utility of our lineage-embedded NSCLC atlas in uncovering biologically meaningful patterns of cell behavior and fate differentiation across tumor, NILT, and PBMC compartments.

NSCLC is generally characterized by a high tumor mutational burden and highly vascularized microenvironment and thus is associated with strong immune infiltration^28,29^. In contrast, ovarian cancer typically exhibits a more immunosuppressive TME, marked by dense stromal barriers and inhibitory factors derived from ascitic fluid^30^. To complement the NSCLC dataset and investigate whether distinct TMEs influence immune clonal architecture, we generated mtscATAC-seq data from primary tumor samples of five ovarian cancer patients, three of whom (SU-O-002, SU-O-004, SU-O-005) had matched peripheral blood samples (**Figure S3A**; **Table S1**). SU-O-005 had undergone neoadjuvant chemotherapy, while other patients were treatment-naive.

In total, we profiled 52,154 immune, malignant, and stromal cells from ovarian tumors and 41,603 PBMCs using mtscATAC-seq (**Figure S3B**). Iterative clustering of the tumor-derived cells identified 40 distinct clusters based on chromatin accessibility profiles, including 13 malignant clusters (*EPCAM*, *KRT18*), 2 stromal clusters (*VWF*, *PECAM1*, *COL1A2*, *FBLN1*), 10 T cell clusters (*CD3D*, *CD8A*, *CD4*), 3 NK/ILC clusters (*NCR1*), 10 myeloid clusters (*CD14*, *LYZ*, *HLA-DQA1*, *KIT*), 1 B cell cluster (*MS4A1*, *PAX5*), and 1 plasma cell cluster (*TNFRSF17*, **Figure S3C**). Annotated stromal and immune cell types comprised cells from multiple donors, whereas malignant clusters were donor-specific, reflecting the interpatient heterogeneity of ovarian tumor subtypes in the dataset (**Figures S3D** and **S3E; Table S1**). Overall, we recovered all major immune cell types previously reported in single-cell studies spanning 24 ovarian tumor samples, supporting the broad cell-type coverage of our dataset (**Figure S3F**)^31,32^. Applying Mitotrek to the ovarian cancer data, we recovered 4,560 clones comprising 20,713 cells (22% of cells passing scATAC QC filters) across donors and tissue sites. Of these, 2,389 clones (53%) contained ≥3 cells, and 287 clones (6%) contained ≥10 cells. The number of clones recovered per patient ranged from 61 to 1,487 (**Figure S3G**). Consistent with our observations in the NSCLC dataset, clone counts were strongly correlated with the number of cells recovered per patient (R = 0.93), and clone assignment rates were uniform across both cell types and tissue sites (**Figures S3H** and **S3I**). All together, the lineage-embedded NSCLC and ovarian tumor datasets provide a robust framework for dissecting lineage dynamics across distinct TMEs.

### Immune clonal landscapes across distinct tissue sites reveals a diverse clonal repertoire of tissue-infiltrating myeloid cells

As mtscATAC-seq enables a measure of clonal diversity across all cell states, we assessed the relative clone sizes of both immune and tumor cells derived from NSCLC and ovarian cancer samples. Specifically, using the Mitotrek, we quantified the distribution of clone fractions by cell type, reasoning that high per-clone fractions reflect cell type-specific clonal expansion events (**Figures 2A** and **S5A**). Overall, tumor cells and adaptive immune cells, including T cells, B cells, and plasma cells, comprised the largest clones observed in both lung and ovarian tumors (**Figures 2B** and **S4A**). Among the 247 clones composed of ≥80% of a single cell type (≥3 cells per clone) detected in lung tumors, defined here as cell type-specific clones, 20.6% were dominated by tumor cells, 11.3% by CD8^+^ T cells, 5.7% by regulatory CD4^+^ T cells (T_reg_), 47.8% by other CD4^+^ T cells, and 5.3% by B/plasma cells. Collectively, these populations comprised over 90% of the cell type-specific clones in lung tumors. In ovarian tumors, of 1,293 cell type-specific clones, 40.9% were dominated by tumor cells, 18.1% by CD8⁺ T cells, and 18.6% by B/plasma cells. Strikingly, only 2.1% and 1.3% were dominated by T_reg_ and other CD4⁺ T cells, respectively, suggesting that clonally expanded CD4⁺ T cells are more prominent in the lung TME than in ovarian tumors.

**Figure 2:**
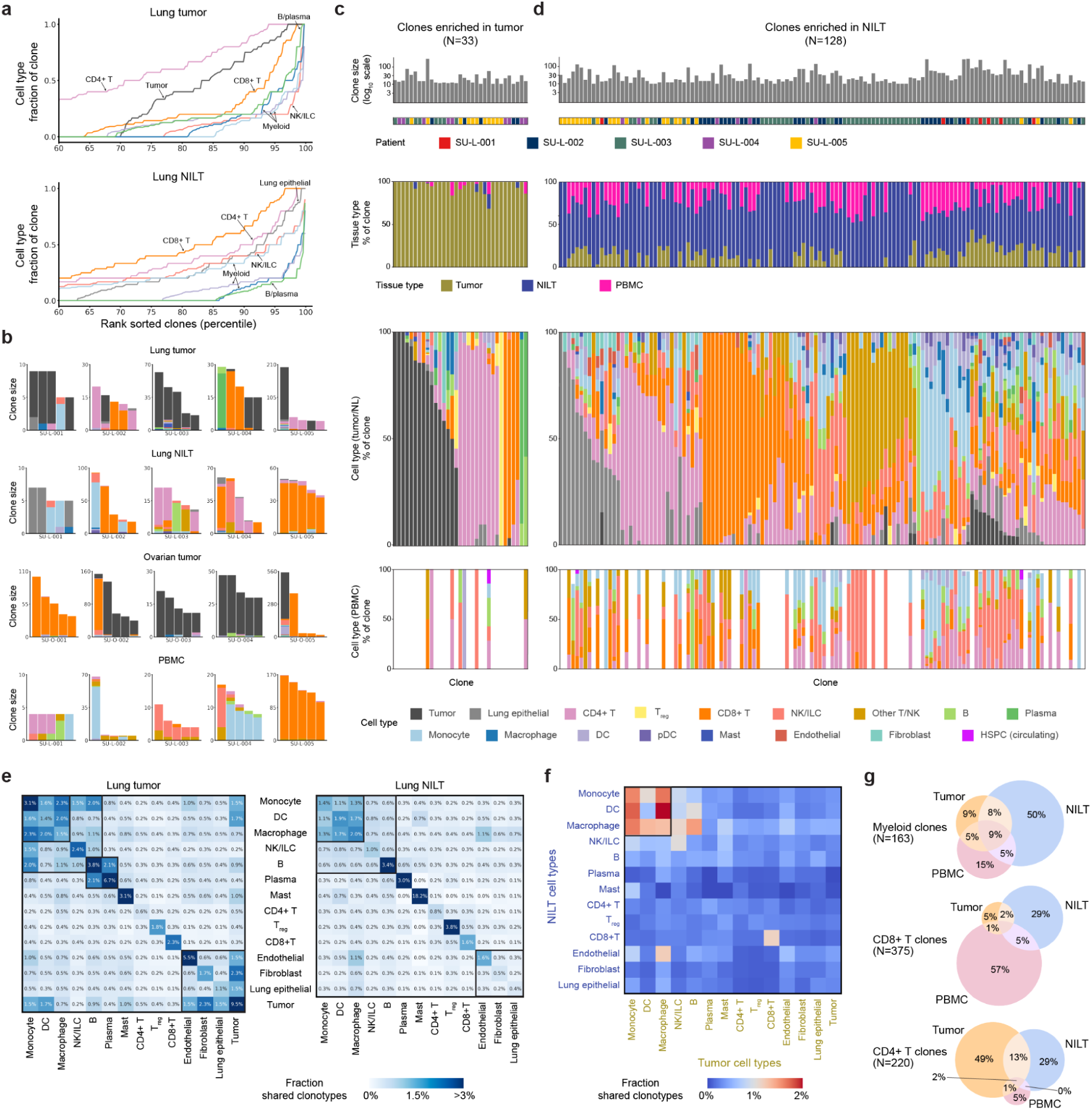
Cross-tissue clonal landscape of tumor-infiltrating immune cells. (A) Cumulative fractions of clones stratified by cell type for cells from lung tumor (top) and NILT (bottom). Clones with ≥5 cells are considered for this analysis. (B) Cell types of single cells belonging to the same clone. The top five most abundant clones with the most common cell type >70% for each patient in the indicated sample are shown. Each bar is colored by cell types of single cells within the clone. (C and D) Clones enriched in tumor (**C**) and in NILT (**D**), as determined by p-value<0.05 from Benjamini-Hochberg adjusted Fisher’s exact test against overall tissue site distribution for clones with at least 10 cells in tumor and NILT. Each column represents a unique clone. (E) Heatmaps showing the fraction of all cell pairs belonging to the same clone and consisting of two cell types within lung tumor (left) and NILT (right). Pairs were restricted to cells from the same donor. (F) Heatmap showing the fraction of all cell pairs belonging to the same clone and consisting of a lung tumor cell type and a NILT cell type. (G) Comparison and overlap of clones (≥3 cells for the indicated cell type) for myeloid, CD8^+^ T, and CD4^+^ T. Myeloid consists of monocyte, macrophage, and DC. N=number of clones considered for each cell type.

Consistent with previous reports of adaptive-like, oligoclonal NK cell responses in human cytomegalovirus infection^20,33–36^, we observed expanded NK cell clones in NILT and peripheral blood. While NK cells were rarely detected in most tumors, one ovarian tumor (SU-O-005) exhibited substantial NK infiltration. These NK cells lacked chromatin accessibility at *CD3D* and *CD8A* loci but showed accessibility at *NCR1* and *GNLY*, confirming their NK identity (**Figure S4D**). Clonal analysis revealed an oligoclonal architecture resembling adaptive lymphocytes (**Figure S4E**). Notably, expanded NK clones were absent from matched peripheral blood, indicating a tumor-restricted NK response (**Figures S4F** and **S4G**). CD8⁺ T cell clonal expansion was also observed in this tumor, suggesting that NK- and T cell-mediated responses can co-occur. Together, these findings highlight distinct clonal expansion patterns across immune cell types and support a role for adaptive-like NK cell responses in human tumors.

The clonal patterns among innate immune cells were similar between NSCLC and ovarian cancer. Myeloid populations, including monocytes, macrophages, and DCs, exhibited smaller clone fractions, compared to adaptive immune cells. Among clones with a substantial number (≥5) of myeloid cells detected in lung tumors, the mean myeloid fraction was 58.5%. In contrast, clones with a substantial number of CD4^+^ T cells, CD8^+^ T cells, and B/plasma cells exhibited higher mean fractions of 78.7%, 78.3%, and 62.1% of the corresponding cell types (**Figure S4B**). Similar patterns were observed in ovarian tumors (**Figure S4C**). These data provide direct evidence that sustained clonal expansion of human myeloid cells following bone marrow egress is limited, compared to lymphoid cell types, which clonally expand following antigen recognition.

We next compared the clonal landscape of lung tumors to the paired normal lung tissue environment. Overall, clone fraction distributions revealed reduced clonal expansion in NILT relative to tumors, as expected (median dominant cell-type fraction per clone 48.3% vs 94.7%, **Figure S5C**). We classified clones with ≥10 cells based on their enrichment in tumor versus NILT (**Figure S5B**). Of 33 tumor-enriched clones, 16 comprised predominantly of tumor cells, collectively accounting for 37.2% of 1,104 cells assigned to these clones. The remaining 17 tumor-enriched clones were dominated by CD4⁺ T cells, CD8⁺ T cells, or B/plasma cells (**Figure 2C**). These adaptive immune clones were largely absent from NILT and PBMCs, with 91.5% of constituent cells detected in tumors, 4.4% in NILT, and 4.1% in PBMCs. In contrast, 128 NILT-enriched clones included a broader range of cell types and had higher representation at other tissue sites (16.9% in tumor and 23.2% in PBMCs, **Figure 2D**). For example, 13 and 15 clones consisted predominantly of invariant NK/T cells and monocytes, respectively, which were minimally observed in tumor-enriched clones. Notably, NILT-enriched clones showed greater cellular heterogeneity (mean dominant cell type fraction: 53.6%) than tumor-enriched clones (85.2%, **Figure S5C**), consistent with reduced clonal expansion in the normal lung tissue environment.

Among the 186 clones that were not preferentially enriched in either the tumor or NILT, 80 are dominated by CD4^+^ T cells and 27 by CD8^+^ T cells (**Figure S5D**). Despite their high cell type purity, these clones were detected in large numbers across tumor, NILT, and peripheral blood (36.7% in NILT, 39.1% in tumor, and 24.2% in PBMC). This observation aligns with previous reports demonstrating concordance between peripheral and intratumoral T cell clone sizes^3,4^. Notably, B cell/plasma cell–dominated clones showed a distinct pattern. While some were restricted to either the tumor or NILT, none spanned multiple compartments including peripheral blood. These results underscore tissue context-specific immune clonal dynamics in NSCLC, suggesting that while T cell clones may exhibit either local or systemic distributions, B/plasma cell clones remain locally confined. Our tumor-NILT enrichment analysis of immune clones implicitly excluded circulating immune cells that may have failed to infiltrate the lung tissues. To identify such clones, we assessed clone enrichment in PBMCs (**Figure S5E**). Among 14 peripheral blood-enriched clones, 10 were dominated by CD8^+^ T cells and 2 by NK cells from a single donor (SU-L-005). Although the majority of cells from these clones were detected in PBMCs (66.0%), smaller fractions were present in tumor (25.3%) and NILT (8.8%), indicating their ability to infiltrate tissue. Together, these results provide a comprehensive view of the clonal composition of immune cells across the human TME and periphery, highlighting cell type- and tissue-specific patterns of immune clonal expansion in human tumors.

### Inter-cell type clonal relationships reveal broad tissue distribution of bone marrow-derived myeloid cells

Next, we systematically quantified lineage relationships and differentiation patterns between cell types within and across tissue sites. We aggregated all clonotypes for each pair of cell types (including self-self pairs) and measured the fraction of clones shared across all distinct cell pairs corresponding to each cell type combination (**Figures 2E** and **S6A; Methods**). We reasoned that two cell types were more related (*i.e.* diverged more recently from the time of sampling) if they frequently shared clonotypes. Across all tissue samples, we identified three broad patterns of cell type lineage relationships: 1) innate immune cells, including monocytes, macrophages, DCs, and NK cells, consistently exhibited high levels of clone sharing, suggesting that they originated from recent hematopoietic output without substantial clonal bottlenecks prior to tissue infiltration, 2) adaptive immune cells and tumor cells showed high levels of intra-cell type clone sharing, reflecting histories of clonal selection and expansion, and 3) stromal cells, including endothelial cells, fibroblasts, and non-tumor epithelial cells, displayed intermediate levels of clone sharing, consistent with their distant embryonic origins relative to HSPC-derived immune cells. We repeated this analysis on PBMCs and observed similar clone-sharing patterns among innate and adaptive immune cells (**Figure S6B**). We noted exceptions to these general trends: non-regulatory CD4^+^ T cells exhibited minimal intra-cell type clone sharing, compared to other adaptive immune cells across tissue sites (0.6%–0.8% in CD4+ T cells, vs. 1.8%–3.8% in T_reg_, 1.6%–2.3% CD8^+^ T cells, 3.4%–3.8% B cells, and 3.0%–6.7% plasma cells). This indicates greater clonal diversity of CD4^+^ T cells despite the clonal expansion of select clones. B and plasma cells exhibited higher levels of clone sharing in lung tumors (2.1%), compared to NILT (0.6%). Endothelial cells demonstrated particularly high intra-cell type clone sharing in lung and ovarian tumors, but not in NILT, consistent with the angiogenic processes indispensable for tumor formation.

We next quantified the lineage relationships between cell types across patient-matched lung tumors, NILT, and/or peripheral blood samples. Among adaptive immune cells, only CD8^+^ T cells consistently exhibited a high degree of clone sharing across tissue sites, suggesting that peripheral expansion during anti-tumor immune response is unique to this population (**Figures 2F** and **S6C**). In contrast, B and plasma cells across different tissues were clonally distinct, supporting the notion that B cell-mediated immunity is locally orchestrated, which has been recently noted in the context of influenza infection^37^. Among myeloid cells, we observed the greatest degree of clone sharing across tissue sites, suggesting that these cells readily infiltrate and differentiate in tissue following bone marrow myelopoiesis and egress. Indeed, myeloid-dominated clones that span multiple tissue sites were the most frequently detected in the NSCLC dataset (27% myeloid clones compared to 8% CD8^+^ T cell clones and 16% CD4^+^ T cell clones, **Figure 2G**). Together, these findings emphasize the distinct clonal dynamics of adaptive and innate immune cells in anti-tumor immunity, highlight the systemic nature of CD8^+^ T cell responses in contrast to the locally restricted nature of B cell responses, and reveal the extensive tissue infiltration and differentiation capacity of bone marrow-derived myeloid cells.

### Intratumoral DC3s are epigenetically and clonally linked to circulating monocytes

It has been reported that specific subpopulations of circulating myeloid cells give rise to tumor-resident mononuclear phagocyte (MNPs) populations that critically shape the TME^13,38,39^. In particular, human DC3s have been identified as a CD1c⁺ dendritic cell subset distinct from cDC2s and to be enriched in NSCLC tumors, and have been suggested to uniquely prime tissue-resident CD8⁺ T cells^27,40–43^. These studies consistently identified a shared expression of monocyte and macrophage gene programs in DC3s in addition to the DC program. However, the epigenetic and ontogenic relationships between human DC3s and other MNPs remain undefined.

Given our observation of broad tissue distribution of bone marrow-derived myeloid cells, we sought to understand the differentiation trajectories of circulating myeloid populations and clonally link them to those in the TME. To achieve higher granularity for myeloid cell type annotation, we first iteratively re-clustered 21,676 myeloid cells from lung tumor, NILT, and peripheral blood in the NSCLC cohort into 13 distinct clusters (**Figures 3A**, **3B**, and **S7A-C**; **Methods**). Monocyte clusters included CD14^+^ classical monocytes and CD16^+^ non-classical monocytes, in both tissues and peripheral blood. Macrophage clusters included alveolar macrophages (AMΦ), interstitial macrophages (IMΦ), and two populations of monocyte-derived macrophages (MoMΦ1 and MoMΦ2), characterized by accessibility at monocyte-associated gene loci, such as *CD14*, *EREG*, and *VCAN*. MoMΦ1 were epigenetically similar to alveolar macrophages, whereas MoMΦ2 were characterized by increased accessibility at *SPP1* and *MERTK*. DC clusters included circulating DCs, cDC1, cDC2, mature DC enriched in regulatory molecules (mregDC)^9,44,45^, and DC3s, recently identified as the dominant DC population in human lung tumors^27^. Given that DCs comprised a minor fraction of myeloid cells in peripheral blood relative to lung tumors and NILTs (4.4% vs. 27.4%, respectively), we were unable to fully resolve DC subtypes in blood and thus grouped circulating DCs as a single population. We validated the myeloid annotations using gene signatures derived from joint single-cell transcriptomic and proteomic profiling of human lung myeloid cells from 35 NSCLC tumors and 29 patient-matched NILTs (**Figures S7D** and **S7E**)^27^. Although prior studies reported subtle transcriptomic differences between tumor- and NILT-derived cells, we observed no corresponding differences in chromatin accessibility, with cells of the same type exhibiting similar epigenetic profiles across tumor and non-tumor tissues (**Figures S7A-E**). Overall, the recovered myeloid cell type profiles were consistent with previous characterizations of human lung tumors and NILTs.

**Figure 3:**
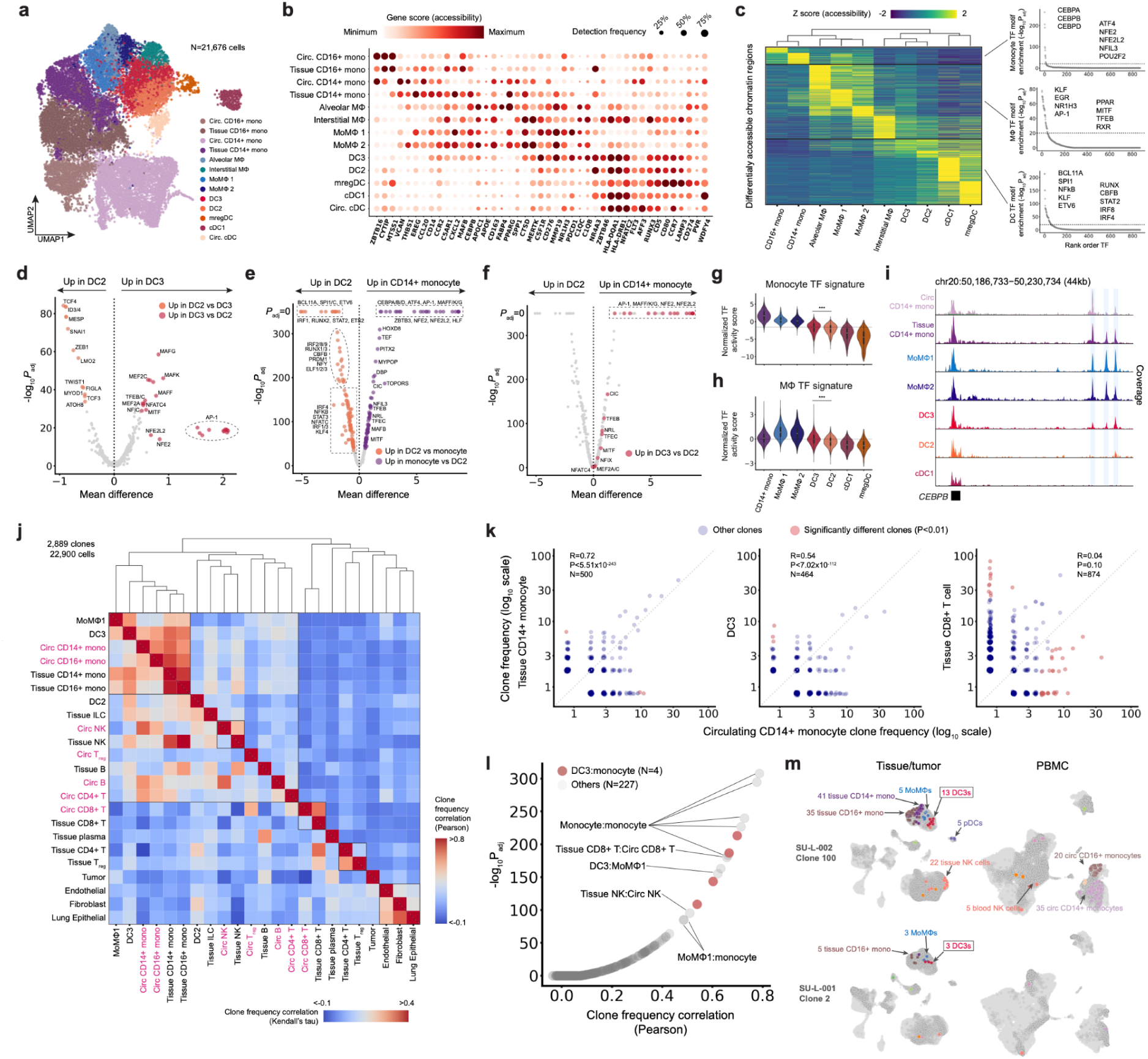
Intratumoral DC3s are epigenetically and clonally related to monocytes. (A) UMAP of myeloid cells from PBMC, lung tumor, and NILT samples of patients with lung adenocarcinoma (LUAD). (B) Column-scaled gene accessibility scores and detection frequencies for the indicated genes. (C) (Left) Heatmap showing marker peaks for all myeloid cell types. The color indicates column-scaled, mean-adjusted number of reads detected in each peak. (Right) Representative transcription factors (TF) motifs that are significantly enriched for monocyte, macrophage, and DC. (D, E, and F) Differentially active TF motifs between DC2 and DC3 (**D**) and DC2 and CD14+ monocytes (**E** and **F**). DC3-up TFs motifs with increased accessibility from (**D**) are highlighted in red in (**F**). P-values are calculated using the Benjamini-Hochberg adjusted Kruskal-Wallis test. (G and H) Average chromVAR motif deviation scores for monocyte and macrophage TF motifs highlighted in (**C**). Kruskal-Wallis test. (I) Chromatin accessibility tracks for the CEBPB locus for the indicated cell types. (J) Cell type-cell type clone frequency correlation across clones (≥5 cells across all samples). Color denotes correlation value, computed using Pearson’s ρ (upper half) and Kendall’s τ (bottom half). Text labels of circulating PBMC cell types are colored pink. (K) Scatterplots comparing clone frequencies of circulating CD14+ monocyte and tissue CD14+ monocyte (left), DC3 (middle), and tissue CD8^+^ T cell (right). Significantly different clones with P<0.05 adjusted Fisher’s exact test are highlighted red. (L) 231 distinct cell type pairs ordered by clone frequency correlation. DC3-monocyte interactions are highlighted in red. (M) representative clones consisting of DC3 and monocytes. For each clone, cell types with at least two cells are highlighted.

Despite extensive transcriptomic and proteomic characterization of lung myeloid cells, the corresponding epigenetic landscape remains understudied. We first used ArchR^46^ to identify differentially accessible regions (DARs) specific to each myeloid cell subtype. Although the subtypes exhibited distinct epigenetic profiles, hierarchical clustering grouped them into three broader categories reflecting their myeloid lineage (**Figure 3C**). IMΦ cells were an exception, clustering with DC subtypes rather than other macrophages; however, due to their rarity across donors (96% of IMΦs originated from a single donor), they were excluded from downstream analyses. To identify key epigenetic regulators defining myeloid identity, we performed transcription factor (TF) motif enrichment analysis on DARs associated with each subtype (**Figures S7F** and **S7G**). In monocytes, CCAAT/enhancer-binding protein (CEBP) motifs were the most differentially active, compared to other MNPs, consistent with prior work establishing CEBPα and CEBPβ as essential for monocyte development^47,48^. These factors are rapidly downregulated during monocyte-to-macrophage differentiation^49^, highlighting their role in maintaining monocyte-specific identity. In macrophages, DARs were strongly enriched for AP-1 family TF motifs (**Figures S7F** and **S7G**), in line with reports showing increased accessibility and 3D chromatin looping at AP-1 binding sites during monocyte-to-macrophage differentiation^50,51^. In addition, motifs associated with other regulators of macrophage differentiation and metabolism, including the retinoid X receptor, MiT-TFE family, liver X receptor alpha, and peroxisome proliferator-activated receptor families, were also more accessible in macrophages^52–59^. In DCs, we observed specific increases in accessibility at binding sites for canonical DC lineage-defining TFs such as BCL11A, IRF4/8, RUNX, CBFB, and ETV6 (**Figures S7F** and **S7G**)^60–62^. Notably, DCs exhibited the highest accessibility of NF-κB family motifs, reflecting NF-κB’s critical role in DC maturation and antigen presentation^63^. Motifs associated with Sp/Krüppel-like factors and early growth response proteins were accessible in both macrophages and DCs but not in monocytes, suggesting these factors promote general monocyte maturation independent of terminal differentiation fate^64–67^.

We next analyzed DARs enriched in DC3s compared to other MNPs. Unlike cDC2s, DC3 DARs exhibited significantly greater accessibility in monocytes and macrophages (**Figures S7H** and **S7I**), suggesting that DC3s may be derived from monocytes and retain monocytic epigenetic features. Indeed, CD14⁺ monocyte DARs were more accessible in DC3s than in other DC subtypes (**Figures 3I** and **S7J**). TF motif analysis of DC3-enriched DARs revealed increased accessibility for motifs associated with monocyte and macrophage TF families, including AP-1, Maf, and MiT-TFE (**Figures 3D-F**). Using chromVAR, we further inferred per-cell TF activity, highlighting concurrent activation of monocyte-, macrophage-, and DC-associated TF programs in DC3s (**Figures 3G**, **3H**, and **S7K**). Notably, CD14⁺ monocytes, but not CD16⁺ monocytes, displayed anticorrelated accessibility of monocyte-versus DC-associated TF motifs (**Figures S7L** and **S7M**), suggestive of transitional states during monocyte-to-DC differentiation in this population. Collectively, these findings demonstrate that DC3s harbor monocytic epigenetic features indicative of a shared ontogeny distinct from classical DCs.

To reconstruct lineage relationships between DC3s and other immune subsets independent of epigenetic profiles, we analyzed clonal frequency correlations across immune subtypes, reasoning that ontogenically related populations are also clonally related. As expected, lineage proximity between circulating and tissue-infiltrating monocytes was recapitulated (**Figure 3J**). Strikingly, DC3s clustered closely with monocytes and MoMΦs (**Figures 3K** and **3L**), whereas cDC2s were less clonally correlated with monocytes (**Figure S7N**). For example, a clone defined by 3068G>A mutation showed 35 CD14^+^ monocytes in the peripheral blood, 41 CD14 monocytes in the tissue, and 13 DCs in the tumor and NILT, likely reflecting continuous infiltration and differentiation (**Figure 3M**). These results provide direct evidence that DC3s are clonally related to circulating monocytes, reinforcing their distinct ontogeny from classical DCs despite phenotypic similarities. Finally, to test whether our findings were lung-specific, we analyzed 8,528 myeloid cells from ovarian tumors, identifying CD14⁺ monocytes, two MoMΦ populations, and two DC populations corresponding to DC3s and cDC2s (**Figure S8A** and **S8B**). Consistent with the findings in NSCLC, ovarian tumor-infiltrating DC3s showed enrichment for AP-1, Maf, MiT-TFE, and other monocyte- and macrophage-associated TF motifs (**Figures S8C-E**), and chromVAR analysis confirmed their distinct epigenetic profile (**Figures S8F-H**). In an ovarian cancer patient with sufficient sampling, intratumoral DC3s were again most clonally related to circulating monocytes (**Figure S8I** and **S8j**), suggesting that the presence of epigenetically monocyte-like DC3s is tumor-agnostic.

### Biased monocyte differentiation fates peripherally reprogram the tumor myeloid compartment

While circulating monocytes have the capacity to differentiate into either macrophages or DCs as they infiltrate inflamed tissue^52,56,62,68,69^, it is not clear whether differentiation fates are entirely dictated by the tissue environment or there is cell-intrinsic bias in the context of human cancer. To address this question, we next asked whether myeloid clones displayed tissue-specific differentiation preferences by comparing their frequencies across lung tumor, NILT, and peripheral blood (**Figures 4A**, **4B**, **S9A**, and **S9B**). Overall, clone frequencies were highly correlated across sites, and the largest clones in tumors and NILT were also detected in blood, indicating infiltration from circulation without tissue site-specific expansion biases. However, within clones, monocytes were proportionally more abundant in NILT, whereas differentiated macrophages and DCs were enriched in tumors (**Figures 4C**, **4D**, and **S9C**). These observations suggest that the TME promotes monocyte differentiation, likely driven by elevated inflammatory cues^69^.

**Figure 4:**
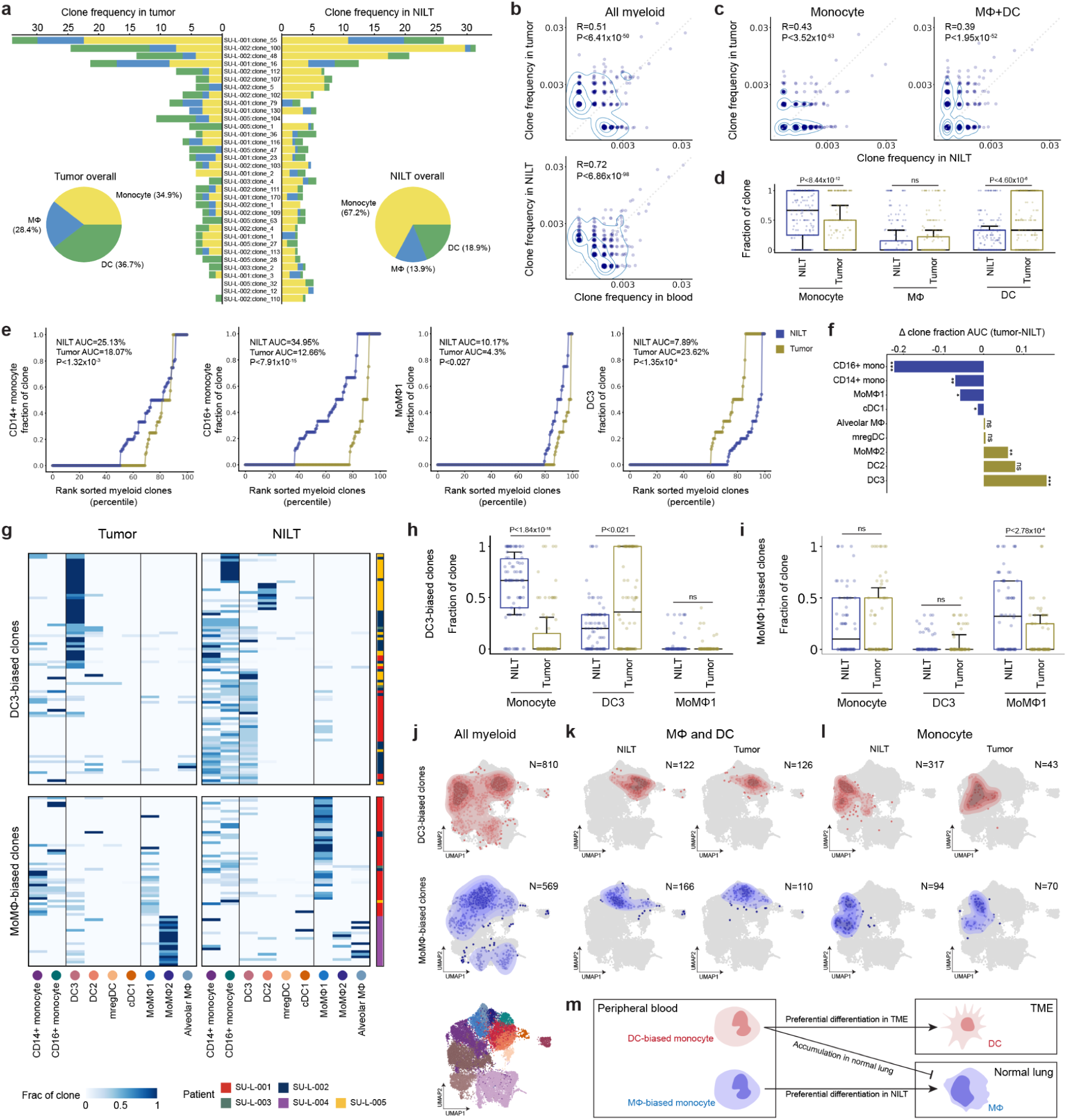
Divergent clonal myeloid differentiation fate in human tissues. (A) Monocyte, macrophage, and DC proportions of largest myeloid clones (≥10 cells), split by tumor and NILT. (B) Scatterplots comparing clone frequencies of circulating myeloid cells with those in tumor (top) and NILT (bottom) myeloid cells. Contours visualize density. (C) Scatterplots comparing clone frequencies of monocytes (left) and macrophage/DC (right) between NILT and tumor. (D) Distribution of the cell type fraction within each myeloid clone (≥5 cells), split by tissue site. Kruskal-Wallis test. (E) Cumulative fraction of clone sizes for the indicated myeloid cell types, split by tissue site. AUC corresponds to the overall clone size for the indicated cell type and tissue site. Kruskal-Wallis test. (F) Summary of AUC differences between tumor and NILT for all myeloid cell types. A positive value indicates the cell type has larger clone sizes in lung tumor compared to NILT. (G) Heatmaps showing cell type proportions split by tumor and NILT for DC-biased (top) and MΦ-biased (bottom) clones. Only clones detected in both tissue sites or clones with at least 3 cells are used to identify cell-type biased clones. (H and I) Distribution of cell type fraction within DC3-biased (**H**) and MoMΦ1-biased (**I**) clones. Kruskal-Wallis test. (J, K and L) Contour plots, representing cell density of DC3-biased clones (top row, red) and MoMΦ-biased clones (bottom row, blue), projected onto the UMAP. All myeloid cells are shown in (**J**), tissue monocytes are shown in (**K**), and MΦ/DC are shown in (**L**). (M) Schematic illustrating divergent clonal differentiation fate for myeloid cells.

To identify the specific myeloid subtypes that monocytes preferentially differentiate into within the TME, we analyzed clone compositions by subtype (**Figures 4E**, **4F**, and **S9D**). Both CD14⁺ and CD16⁺ monocytes were less represented in tumors compared to NILTs, while SPP1⁺ MoMΦ2 and DC3s were preferentially enriched in tumors. Notably, MoMΦ1 was more abundant in NILTs. These clone-level observations align with prior donor-matched NSCLC studies reporting, in overall fractions, monocyte enrichment in NILTs, DC3 enrichment in tumors, and variable enrichment of MoMΦs depending on their specific subtypes^8,27,45,70^.

If differentiation were purely driven by the tissue environment, the same clone would display different myeloid subtype compositions between tumor and NILT. To test this, we clustered clone-level subtype compositions stratified by tissue site (**Figure 4G**). Interestingly, we observed clone-intrinsic lineage biases, with clones consistently favoring differentiation toward either DCs or macrophages, independent of tissue site. DC-biased clones were enriched for DC3s in tumors and consisted primarily of monocytes in NILTs, with minimal macrophage representation across sites. In contrast, macrophage-biased clones exhibited distinct behaviors depending on the MoMΦ subtype: MoMΦ1-biased clones were depleted of monocytes and enriched for MoMΦ1 cells in NILTs (**Figure 4I**), whereas MoMΦ2-biased clones almost exclusively consisted of MoMΦ2 cells in tumors. These findings suggest that monocytes possess intrinsic biases in both differentiation fate and site preference.

We next grouped myeloid cells derived from DC-biased and macrophage-biased clones to compare their epigenetic profiles. Across tissue sites, cells from the two groups clustered distinctly based on epigenetic profile, even within the same myeloid subtype such as CD14^+^ monocytes in both tissue and periphery (**Figures 4J-l**). Remarkably, even prior to tissue infiltration, circulating CD14⁺ monocytes from the two groups were epigenetically distinct in peripheral blood, with monocytes within each group showing greater similarity to one another than to those in the other group (**Figures S9E** and **S9F**). Analysis of TF motif accessibility revealed functional distinctions between the CD14⁺ monocytes of these groups. DC-biased monocytes exhibited enrichment of proinflammatory regulatory programs, while macrophage-biased monocytes showed enrichment of immunosuppressive programs. In the peripheral blood, motifs for NFκB family members, IRF family members, STAT2, and BLIMP-1 were differentially accessible in DC-biased monocytes, consistent with their known roles in regulating innate immune responses, antiviral defense, and antigen presentation^71,72^ (**Figures S9G**). Upon tumor infiltration, IRF family and BLIMP-1 motifs remained differentially accessible in DC-biased monocytes, while ID3/ID4 motifs additionally became selectively accessible in this population, further supporting their antitumor potential^73^ (**Figures S9H**). Conversely, circulating macrophage-biased monocytes displayed increased accessibility of Maf family motifs, critical regulators of macrophage differentiation, along with motifs associated with immunosuppressive TFs such as NRF2 and RUNX1/2^74–77^.

Because motif enrichment alone does not fully resolve TF identity among homologous family members, we further analyzed gene body accessibility of TFs across monocyte populations (**Figures S9I** and **S9J**), reasoning that functionally active TFs are more likely to exhibit both motif and gene body accessibility. We found that *IRF1*, *IRF3*, and *BLIMP-1* were differentially accessible in DC-biased monocytes in both peripheral blood and tissue. *STAT2* was only differentially accessible in circulating DC-biased monocytes, while *IRF7* was only differentially accessible in tissue-infiltrating DC-biased monocytes, suggesting their roles are upstream and downstream of tumor infiltration, respectively. In macrophage-biased monocytes, *NFE2* and *MAF* (encoding cMaf) were differentially accessible in circulation. Notably, cMaf is a key transcriptional regulator of macrophage immunosuppressive phenotypes, promoting *IL10* expression and suppressing proinflammatory *IL12* through direct binding of cis-regulatory elements^78–80^. Taken together, these findings suggest that circulating monocytes exhibit intrinsic epigenetic biases that predispose them to differentiate into either DCs or macrophages within the TME, contributing to their distinct functional roles in shaping the antitumor innate immune response.

## Discussion

Here, we performed mtscATAC-seq on 218,715 cells across 23 matched tumor, non-involved tissue, and peripheral blood samples from patients with lung and ovarian cancers to generate a clonally resolved map of the human innate immune response to cancer. To ensure the accuracy of clonal tracing, we developed Mitotrek, a tailored analysis framework that prioritizes clonal assignment fidelity (over sensitivity or resolution), which enabled the delineation of clonal relatedness across and between cell types in solid tumors and tissues. By applying this approach to patient-matched samples from multiple tissue types, we simultaneous profiled epigenetic states and clonal relationships of tens of thousands of immune cells per patient, while mitigating technical and biological sources of false-positive lineage connections. The validity of this approach was demonstrated using a published dataset with ground-truth lineage information, enabling robust characterization of the clonal landscape of tumor-infiltrating innate immune cells in human samples.

Using Mitotrek, we assessed the clonal diversity of immune cell types across tumor, non-tumor tissue, and peripheral blood, revealing distinct patterns of expansion between innate and adaptive compartments. Innate immune cells in tumors consistently exhibited high clonal diversity without evidence of tissue-specific selection, suggesting they are primarily derived from a diverse pool of circulating precursors and do not clonally expand in the TME. In contrast, adaptive immune cells and tumor cells exhibited the highest levels of clonal expansion, with the largest clones accounting for the majority of these populations, consistent with the role of antigen-driven selection in shaping the adaptive immune repertoire. Clonotype sharing across tissue sites further revealed that myeloid populations were highly related, supporting a model in which tissue myeloid cells are replenished by a common reservoir of bone marrow-derived precursors circulating through the body. Epigenetic profiling of myeloid populations in human lung tissue and peripheral blood provided insights into the regulatory landscape of DC3s. Compared to classical dendritic cells, DC3s exhibited increased chromatin accessibility at monocyte-associated regulatory elements, implicating transcription factors central to monocyte and macrophage biology. These findings corroborate previous transcriptomic and proteomic studies linking DC3s to monocytes^27,40,41,43^. Clone frequency correlation analysis revealed that DC3s, but not cDC2s, were clonally related to monocytes in both circulation and tissue. A recent study in mice showed that DC3s arise from bone marrow progenitors shared with monocytes^81^. While the clonal resolution of our dataset cannot rule out this possibility, the relative rarity of DC3s in peripheral blood and their abundance in tumor lesions strongly suggest that human DC3s in tumors differentiate directly from monocytes upon tissue infiltration.

A key finding of our work is that phenotypically diverse tumor-infiltrating myeloid cells were clonally related to distinct subsets of circulating monocytes. Importantly, clones were consistently biased toward either DC or macrophage fates in both tumor and non-involved tissue. Both DC-biased and macrophage-biased myeloid clones contained monocytes, suggesting that this bifurcation in differentiation fate had occurred downstream of common dendritic progenitor-granulocyte myeloid progenitor bifurcation in bone marrow myelopoiesis. These differentiation biases were accompanied by distinct chromatin accessibility patterns, detectable even in peripheral blood monocytes prior to tissue infiltration. Specifically, macrophage bias was linked to increased accessibility of AP-1 family transcription factors, c-Maf, and NRF2. Interestingly, a recent study identified a macrophage subset defined by FOSL2 (an AP-1 factor) activity and derived from circulating monocytes as strongly associated with glioma malignancy^38^, supporting our observation that AP-1–enriched monocytes are predisposed to macrophage differentiation within the tumor microenvironment. Our results suggest that the tumor myeloid compartment may be peripherally programmed, posing an alternative to the view that intratumoral monocyte differentiation is primarily driven by local environmental cues.

The DC- or macrophage-bias observed in circulating monocytes may reflect systemic reprogramming of myelopoiesis by tumor-derived signals. Tumors are known to secrete cytokines that act distally on HSPCs, altering the output and fate of bone marrow-derived myeloid cells^82^. Supporting this, DC-biased monocytes exhibited increased chromatin accessibility at IRF, STAT2, and NF-κB motifs, suggesting priming by type I interferons, TNF-α, and IL-1 signaling^83,84^. These findings imply that the tumor may enable epigenetically pre-condition monocytes in the periphery, shaping their differentiation trajectory before tissue infiltration and contributing to the observed fate biases within the TME.

Several limitations of our study should be addressed in future work. First, our findings are based on a relatively small cohort of patients with lung and ovarian cancers. Although we deeply profiled each patient and observed consistent patterns across patients and cancer types, validation in larger cohorts and additional tumor types will be important to establish the generalizability of our conclusions. Second, our approach does not capture the temporal dynamics of monocyte infiltration and differentiation. Longitudinal sampling or integration with temporally resolved data will be necessary to definitively establish the sequence of events that shape the tumor myeloid compartment. Lastly, our current approach collapses subclonal relationships. Future strategies with the ability to resolve subclonal architectures would provide a clearer view and stronger evidence for the differentiation dynamics we propose.

In summary, our study provides a high-resolution clonal and epigenetic atlas of the human tumor myeloid compartment, offering new insights into the fundamental biology of innate immunity and potential avenues for therapeutic intervention. The clonal differentiation fate biases we observed in tumor-infiltrating monocytes have important implications for myeloid-targeted immunotherapies. DC3s and tumor-associated macrophages may represent attractive targets for augmenting antitumor immunity, as they appear to be pre-programmed for proinflammatory and immunosuppressive functions, respectively. Strategies that selectively deplete monocytes biased toward a macrophage fate while preserving those poised for dendritic cell differentiation may tip the balance toward a DC3-rich tumor microenvironment, potentially enhancing tumor antigen presentation and T cell priming. More broadly, our results highlight the power of mitochondrial lineage tracing for dissecting the clonal dynamics of immune cells in human cancers and reveal new opportunities for harnessing the innate immune system in cancer immunotherapy.

## Methods

### Human subjects

Fresh ovarian tumors, lung tumors, NILT samples, and peripheral blood were collected at the time of surgery by Stanford Tissue Procurement Shared Resource facility with the appropriate written informed consent and institutional IRB approval. Summary statistics and patient history are available in Supplementary Table 1. For the early-stage NSCLC dataset, exclusion criteria included previous systemic treatment or radiotherapy. For the ovarian cancer dataset, exclusion criteria included tumors of non-ovary or unknown origin.

### Tissue processing

All tumor and adjacent healthy tissues were procured following surgical resection. Samples were dissociated and viably cryopreserved for downstream library preparation and sequencing. In brief, solid tumor specimens on ice were minced to pieces of <1 mm3 and transferred to 5 ml of digestion medium containing DNase I (100 µg/ml) and collagenase P (2 mg/ml) in Advanced DMEM/F-12. Minced tissue was transferred into C-tubes for use in the gentleMACS Octo Dissociator system at 37 °C at 20 rpm for 20 min. After digestion, the cell suspension was filtered through a 70-μm filter, which was washed with an additional 10 ml of DMEM/F-12 and centrifuged the sample at 400g at 4°C for 5 min. Any residual undigested tissue was further digested for an additional 20-min incubation with additional digestion medium. After centrifugation, the supernatant was discarded, and the pellet was resuspended in 500 µl of ACK red blood cell lysis buffer and incubated for 1 min on ice, followed by the addition of ice-cold PBS. The cell count and viability were determined by trypan blue staining by using a Countess II FL automated cell counter, before proceeding to cell-sorting.

### Fluorescence-Activated Cell Sorting (FACS)

Cells were classified into peri-tumoral T cells (CD45⁺CD3⁺), other peri-tumoral lymphocytes (CD45⁺CD3⁻), and malignant or stromal cells (CD45⁻CD3⁻). The antibodies used included anti-human CD45 conjugated to V500 (clone HI30, 560779, lot 7172744, BD Biosciences) and anti-human CD3 conjugated to fluorescein isothiocyanate (FITC) (clone OKT3, 11-0037-41, lot 2007722, Invitrogen), both diluted at 1:200. Live/dead staining was performed using propidium iodide (P3566, Invitrogen) at a final concentration of 2.5 μg/mL. Cell sorting was conducted on a BD FACSAria™ III cell sorter (BD Biosciences).

### Preparation of mtscATAC-seq libraries

For the generation of mtscATAC-seq libraries, we adapted the 10× Genomics scATAC-seq platform NextGEM v1.1 kits. In brief, mtscATAC-seq was performed with modifications to the “Nuclei Isolation for Single Cell ATAC Sequencing” (CG000169 Rev D) user guide, where we fix and permeabilize cells to retain mitochondria and mtDNA within their host cell by removing Tween 20 as part of the lysis buffer.^19^ For the library preparation we follow the “Chromium Next GEM Single Cell ATAC Reagent Kits v1.1” (CG000209 Rev F), user guide with only minor modifications as described and highlighted below and otherwise refer the reader to the original and highly detailed workflow by 10× Genomics^19^.

### Sequencing and upstream processing of mtscATAC-seq data

All libraries were sequenced on an Illumina Novaseq 6000 device using a 10×16×151×151 read configuration to accommodate the 10× ATAC cell barcode in the i5 piece of the read. Libraries were sequenced to a target of 30,000–35,000 reads/cell as previously recommended^19^. Raw .bcl files were converted into per-sample .fastq files using Illumina bcl2fastq.

Initial processing of mtscATAC-seq data was performed using the CellRanger-ATAC Pipeline v.2.0.0 by mapping scATAC-seq reads with cellranger-atac count to the GRCh38 reference genome, hardmasked for regions that would otherwise interfere with mapping to the mitochondrial genome (as previously detailed^17^). The outputs included fragments files for downstream epigenomics analyses and .bam files for mtDNA genotyping.

Mitochondrial genotyping was performed on these fragment files with mgatk in tenx mode using the barcodes identified as cells by CellRanger (*i.e.* cells passing ATAC filter). Only cells with at least 10× coverage of the mitochondrial genome were included in the analysis, achieving a median coverage of 30×-70× across experiments.

### scATAC-seq QC, dimensionality reduction and clustering

CellRanger output fragment files were loaded and converted to Arrow files using the createArrowFiles function in ArchR. Quality control metrics were computed for each cell, and only cells with TSS enrichments greater than 4 were kept for all samples. Cells were also filtered based on the number of unique fragments sequenced using a cut-off of 1000. Doublet scores for all cells were computed using the ArchR functions addDoubletScores with k=10, knnMethod=”LSI”, and LSIMethod=1.

Sample groupings were defined based on cell types, tissue sites, and disease status (eg. all cells, lung tumor and NILT samples from patients with NSCLC in **Figure 1**; myeloid cells, all samples from patients with NSCLC in **Figure 3**). An ArchR project was then created for each of the sample groupings and doublets were filtered with filterDoublets with a filter ratio of 1. For each ArchR project, dimensionality reduction was performed with addIterativeLSI using default parameters to embed ATAC-data in latent semantic indexing (LSI) space. Next, clustering was performed with addClusters using default parameters.

### Annotation of mtscATAC-seq dataset

An iterative clustering approach was used to annotate cells, where after each round of clustering, select clusters with relatively high epigenetic similarity (eg. T and NK cells) are merged and reclustered to achieve desired granularity and higher clustering accuracy. For the NSCLC lung tumor/NILT data sample grouping and ovarian tumor sample grouping, clusters were annotated based on gene score of known marker genes, including *EPCAM*, *KRT18* (epithelial and tumor cells), *VWF*, *PECAM1* (endothelial cells), *COL1A2*, *FBLN1* (fibroblasts), *CD3D*, *CD4*, *FOXP3*, *CD8A*, *KLRD1*, *NCR1* (T and NK cells), *MS4A1*, *PAX5* (B cells), *TNFRSF17*, *VOPP1* (plasma cells), *CD14*, *LYZ* (monocytes), *APOC1*, *CD163* (macrophages), *HLA-DQA1*, *ZBTB46* (DCs), *CLEC4C* (PDCs), *TPSAB1*, *KIT* (mast cells).

9 initial clusters in the lung tumor/NILT data annotated as epithelial or tumor cells were grouped and reclustered, resulting in 22 clusters. 9 clusters were annotated as tumor for satisfying the following criteria 1) highly patient specific and 2) enriched over four-fold in tumor compared to NILT. The other 13 clusters were annotated as lung epithelial subtypes based on high gene score of following signatures *AGER*, *PDPN*, *CLIC5* (alveolar type 1), *SFTPB*, *SFTPC*, *SPTPD*, *MUC1*, *ETV5* (alveolar type 2), *FOXJ1*, *TUBB1*, *TP73*, *CCDC78* (ciliated), *KRT5*, *KRT17*, *MIR205HG* (basal), *MUC5B*, *SCGB1A1*, *BPIFB1*, *PIGR*, *SCGB3A1* (secretory).

6 initial clusters in the lung tumor/NILT data annotated as T/NK cells were further divided into 19 clusters. 5 initial clusters in the ovarian tumor data annotated as T/NK cells were further divided into 13 clusters. These T/NK clusters were annotated using known marker genes, including *CD3D* (broad T), *CD8A* (CD8^+^ T), *CD4* (CD4^+^ T), *GNLY*, *NCR1* (NK/ILC), *FOXP3* (T_reg_). To support the marker gene-based annotation, cells were projected to human PBMC reference data using the Azimuth application ^85^. The Azimuth-predicted cell types were dominated by CD4^+^ T, CD8^+^ T, T_reg_, NK, and ILC, which are all associated with distinct clusters and consistent with the mark-gene based annotations. One cluster in the lung data is marked by the gene score pattern of *CD3D*+, *CD8A*-, *CD4*-, *CD56*+ and high gene score of NK signature (*GNLY*, *PRF1*, *GZMB*, *KLRB1*, *CCL3*, *KLRF1*, *NCR1*), which was defined as invariant NKT cells.

### Gene signature scoring

Gene scores for individual genes are computed as implemented in ArchR. When computing the composite gene score for a gene signature, gene scores of individual genes in the signature were z-score normalized across all cells, and each cell is then scored by taking the mean z-scaled gene scores in the gene signature.

### Peak calling and motif analysis

For the NSCLC myeloid epigenetic analysis, peak calling was performed as implemented in ArchR. Both level1 (monocyte, macrophage, DC) and level 2 (eg. CD14+ monocyte, DC3, MoMΦ1, etc) cell type annotation was used as grouping in addReproduciblePeakSet() to identify regulatory elements associated with both broad myeloid cell types as well as subtypes. Differential peak analysis was performed using getMarkerFeatures. TF motifs enriched in peak sets are identified using peakAnnoEnrichment. Per-cell TF motif activities were calculated using chromVAR^86^.

### Mitochondrial genotyping and clone calling

By default, mgatk calls high-confidence heteroplasmic variants by selecting variants with strand correlation>0.65 and variance-mean ratio>0.01. In this study, most donors have multiple samples from different tissue sites. Given that the default mgatk filters are already conservative, an union approach to selecting high-confidence heteroplasmic variants is employed to increase sensitivity, where if a variant passes the default mgatk filters in any sample, then it is considered a valid, informative variant in all other samples from the same donor (eg. if a variant is determined to be high quality in the lung tumor sample, it is unlikely to be spurious if it is also observed in the peripheral blood sample from the same donor). We noted one polymorphic and highly homologous region (CCCTCCC in GRCh38 chrM:307-314) that led to spurious connections under the relaxed filtering criteria and explicitly disregarded variants from this region. This mitochondrial variant processing procedure for clone calling outputs a cell by variant heteroplasmy matrix combining all samples for each donor and is implemented in the mitotrek.processing module.

Clone is then called for each donor using mitotrek.core.assign_cell_to_clones. Variants present in >20% cells from a donor (likely technical or homoplasmic) or less than 3 cells are removed. The heteroplasmy matrix is then binarized with a cutoff of 0.07 based on the rationale that exact heteroplasmy levels are not reliable given the stochasticity from mitochondrial genome distribution during cell division and variance in per-cell mitochondrial genome coverage. The heteroplasmy cutoff value is chosen based on benchmarking experiments using published data (**Supplemental Figure 2g-h**).

To convert mitochondrial variants to clones, an undirected weighted graph is constructed in which vertices are variants and edges are defined by the Pearson correlation coefficient between two variants across cells, computed from the binarized heteroplasmy matrix. After removing edges with weights less than 0.5, each connected component in the graph was treated as a distinct clonotype. Most resulting connected components contained only one variant, whereas highly correlated variants (ie. frequently co-occuring in cells) were grouped into one connected component. For clones defined by a single variant, any cell positive for the associated variant in the binarized heteroplasmy matrix is assigned to the clone. For clones defined by multiple variants, a cell is required to be positive for all associated variants. Finally, cells assigned to multiple clones are discarded.

### Benchmarking Mitotrek clone assignment

Full-length scRNA-seq via the Smart-seq2 technology of single-cell derived colonies from two donors published by Ludwig et al.^16^ was downloaded as .fastq data from the Gene Expression Omnibus (accession GSE115214). Mitochondrial genotyping was performed using mgatk and cells with at least 100× mitochondrial genome coverage were retained for downstream analysis. Clones were called using mitotrek.core.assign_cell_to_clones with a binarization cutoff threshold set from 0.01 to 0.15. For each ground truth clone label (established in the experimental protocol by physical separation of the clones), the predicted clone label was determined by taking the mode of called clones among cells in the ground truth clone to compute accuracy, which is implemented in mitotrek.core.clone_calling_accuracy.

### Clone sharing analysis

We consider two mutually exclusive sets of cells *A* and *B* (eg. two cell types). There are N clones *c*_1_, *c*_2_, …, *c*_*N*_ containing at least one cell in *A*∪*B*. Note that each clone *c_i_* belongs to a set *P_j_* which contains all clones detected in donor *j*. We summarize the clone counts in the two groups as a *N*×2 matrix *X* where *X*_*i*,1_ and *X*_*i*,2_ correspond to number of clone *c* cells in *A* and *B*, respectively. The fraction of clones shared between the two groups is computed as

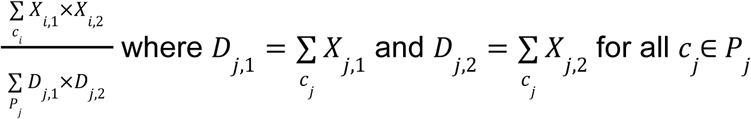

The numerator computes the number of observed cell pairs between two groups that share a clone. The denominator is a normalization factor that represents all possible cell pairs between two groups, accounting for only cell pairs within the same donor, since cross-donor cell pairs are not valid.

### Quantification and statistical analysis

Statistical analysis of single-cell sequencing data was performed in python (v3.9.4) and R (v4.0.5). Statistical analysis of flow cytometry data was performed in GraphPad Prism (v9.0)

## Code Availability

The chromVAR analysis software for epigenetic analysis of scATAC-seq data is available on Github (https://github.com/GreenleafLab/ArchR). The mgatk software for processing sequencing data for single-cell mitochondrial variant calling is available on Github (https://github.com/caleblareau/mgatk). The Mitotrek software developed in this work for clone calling using single-cell mitochondrial variant data is available on Github (https://github.com/vincent6liu/mitotrek). Any additional custom code used for computational data processing and analysis are available from the authors upon request.

## Data Availability

Raw sequencing and processed chromatin accessibility and mtDNA mutation calls are available at the Gene Expression Omnibus accession **GSE302113** with reviewer access token **udwbeuumxvcrpod**.

---

**Supplemental Figure 1:**
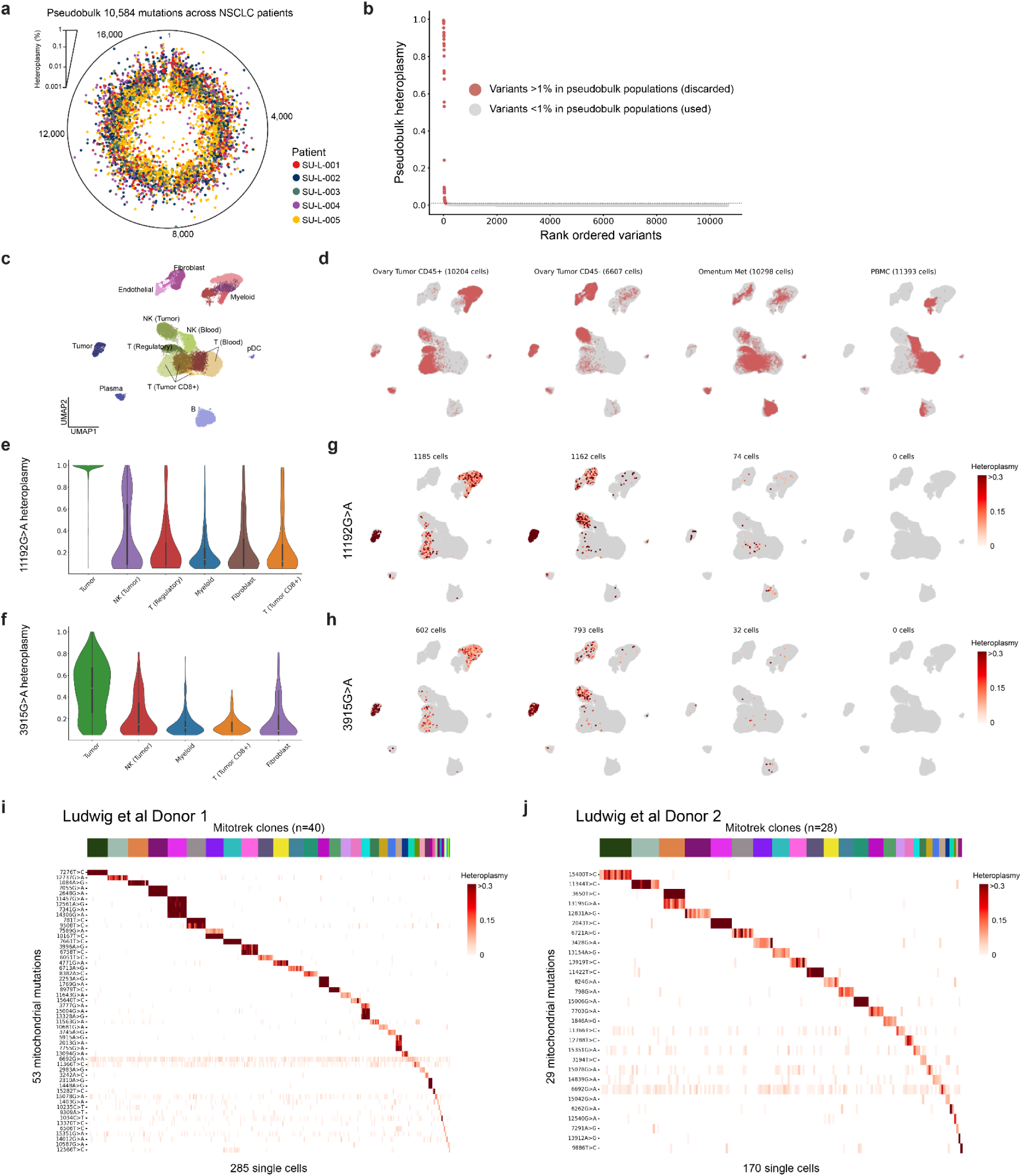
Benchmarking Mitotrek using ground-truth data. (A) Distribution of mgatk-nominated variants along the mitochondrial genome, averaged across cells and colored by patient. (B) Pseuboulk heteroplasmy for all variants detected by mgatk in across all samples. Variants with >1% pseudobulk heteroplasmy are excluded from downstream analysis. 1% is chosen as the conservative threshold after observing the overall heteroplasmy distribution. (C) UMAP embeddings of tumor-infiltrating immune cells from matched primary (ovarian) and metastatic (omentum) tumors, and PBMCs from HGSC patient SU-O-005. (D) Distribution on the UMAP of cells from indicated samples. Cells from the primary ovarian tumor were sorted by CD45 to separate tumor-infiltrating immune cells. (E and F) Heteroplasmy levels of the indicated tumor-specific mitochondrial variants in tumor-infiltrating immune cells processed together with tumor cells. (G and H) Mutations projected onto UMAP embeddings across samples. Tumor-specific variants are indiscriminately detected at lower heteroplasmy levels in all cells from the same sample, suggesting mitochondrial transfer and/or technical artifacts (ambient mtDNA). (I and J) Heatmap showing the heteroplasmy levels of variants (rows) that are identified as clone markers to group cells (columns) in each Mitotrek clone. Position of each variant and the base pair change are shown.

**Supplemental Figure 2:**
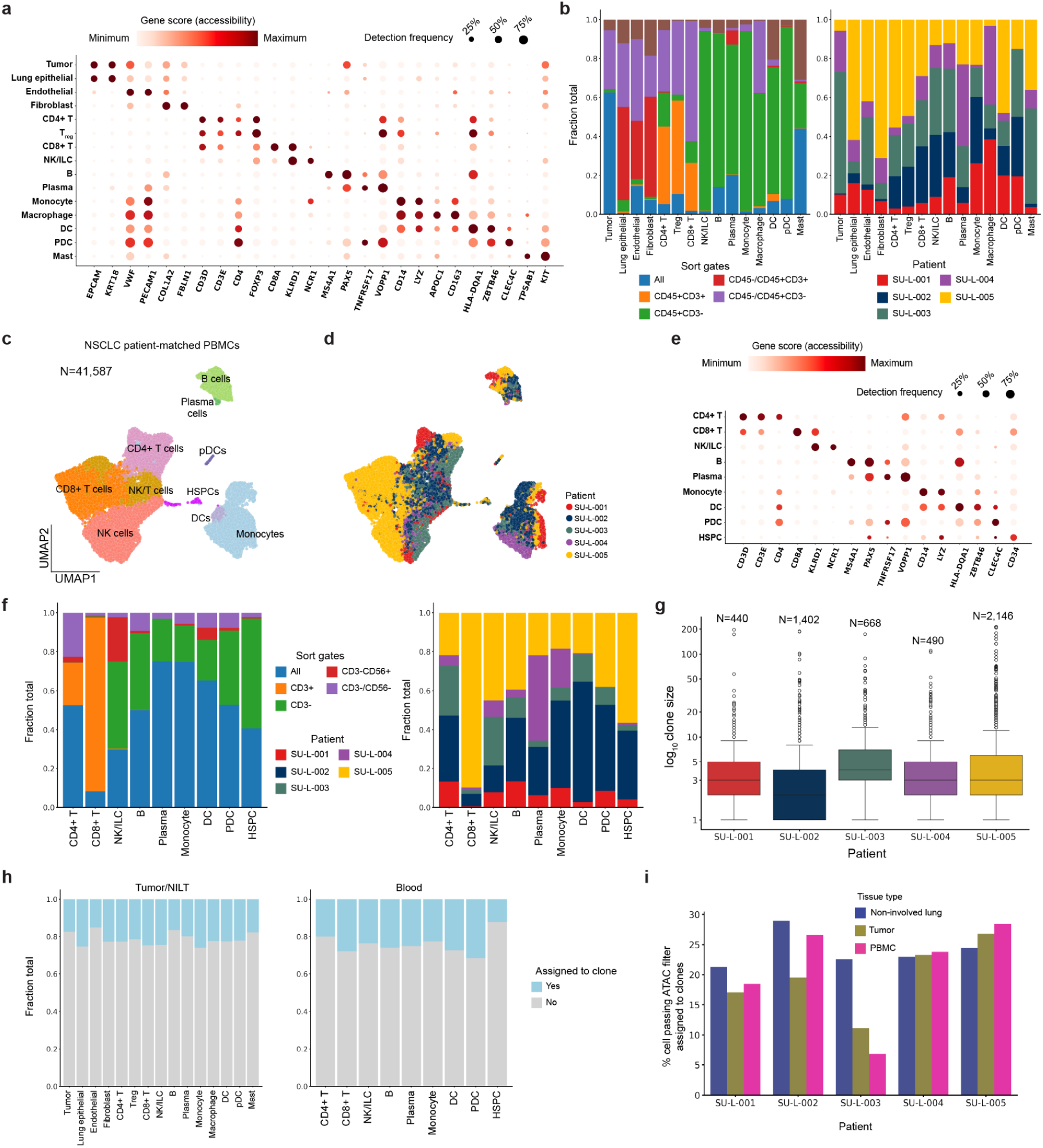
NSCLC data summary and cell type annotation. (A) For cell types annotated in tumor and NILT samples, column-scaled gene accessibility scores and detection frequencies for the indicated genes. (B) For tumor and NILT samples, bar plots indicating (left) relative proportions of markers used for sorting that were detected in each cell type (certain samples were not sorted, and select sorted samples were merged during single-cell capture, due to sample-specific considerations to optimize single-cell yield) and (right) relative proportions of cells from each patient detected in each cell type. (C) UMAP of 41,587 PBMCs from patients with lung tumors. (D) UMAP of cells colored by patient identity. (E) For cell types annotated in PBMC samples, column-scaled gene accessibility scores and detection frequencies for the indicated genes. (F) For PBMC samples, bar plots indicating (left) relative proportions of markers used for sorting that were detected in each cell type (certain samples were not sorted, and select samples were additionally sorted using CD56 to enrich innate lymphocytes and myeloid cells) and (right) relative proportions of cells from each patient detected in each cell type. (G) Clone size distributions for patients with NSCLC. (H) Bar plots summarizing relative proportions of cells assigned to clones across cell types in tumor/NILT and blood. No significant cell-type bias was observed. (I) Fraction of cells passing ATAC filters that are successfully assigned to clones.

**Supplemental Figure 3:**
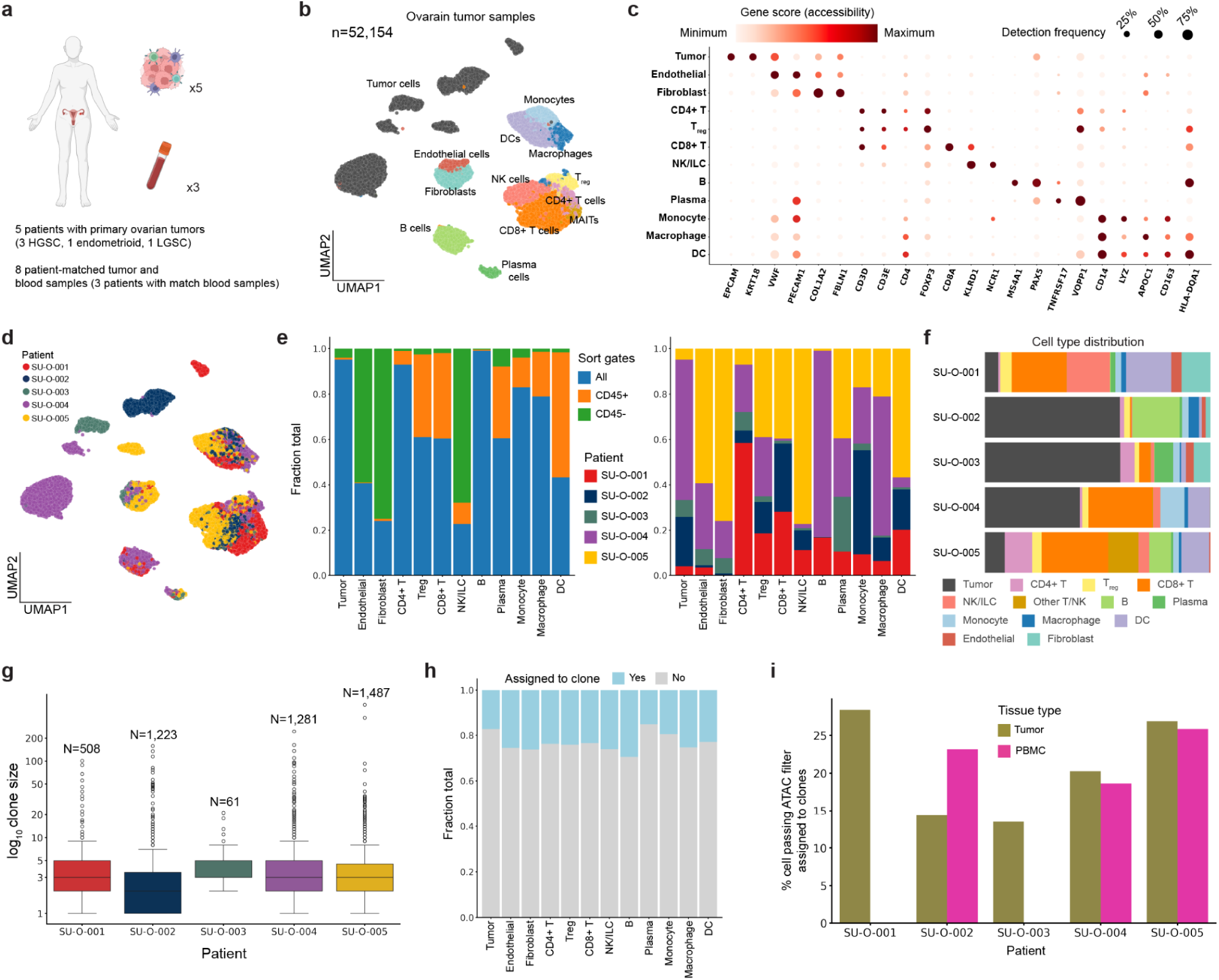
Ovarian cancer data summary and cell type annotation. (A) Schematic summarizing patient and sample information for the ovarian tumor data. (B) UMAP of 52,154 cells in ovarian tumors. Cell types denoted by color are inferred after iterative sub-clustering of each of the myeloid, lymphoid, and stromal compartments. C) For cell types annotated in ovarian tumor samples, column-scaled gene accessibility scores and detection frequencies for the indicated genes. (D) UMAP of cells colored by patient identity. (E) For ovarian tumor samples, bar plots indicating (left) relative proportions of markers used for sorting that were detected in each cell type (certain samples were not sorted, and select sorted samples were merged during single-cell capture, due to sample-specific considerations to optimize single-cell yield) and (right) relative proportions of cells from each patient detected in each cell type. (F) Normalized bar plot showing cell type composition for each patient. (G) Clone size distributions for patients with ovarian cancer. (H) Bar plots summarizing relative proportions of cells assigned to clones across cell types in ovarian and blood. No significant cell-type bias was observed. (I) Fraction of cells passing ATAC filters that are successfully assigned to clones. Peripheral blood samples were obtained from three patients. Created with BioRender.com.

**Supplemental Figure 4:**
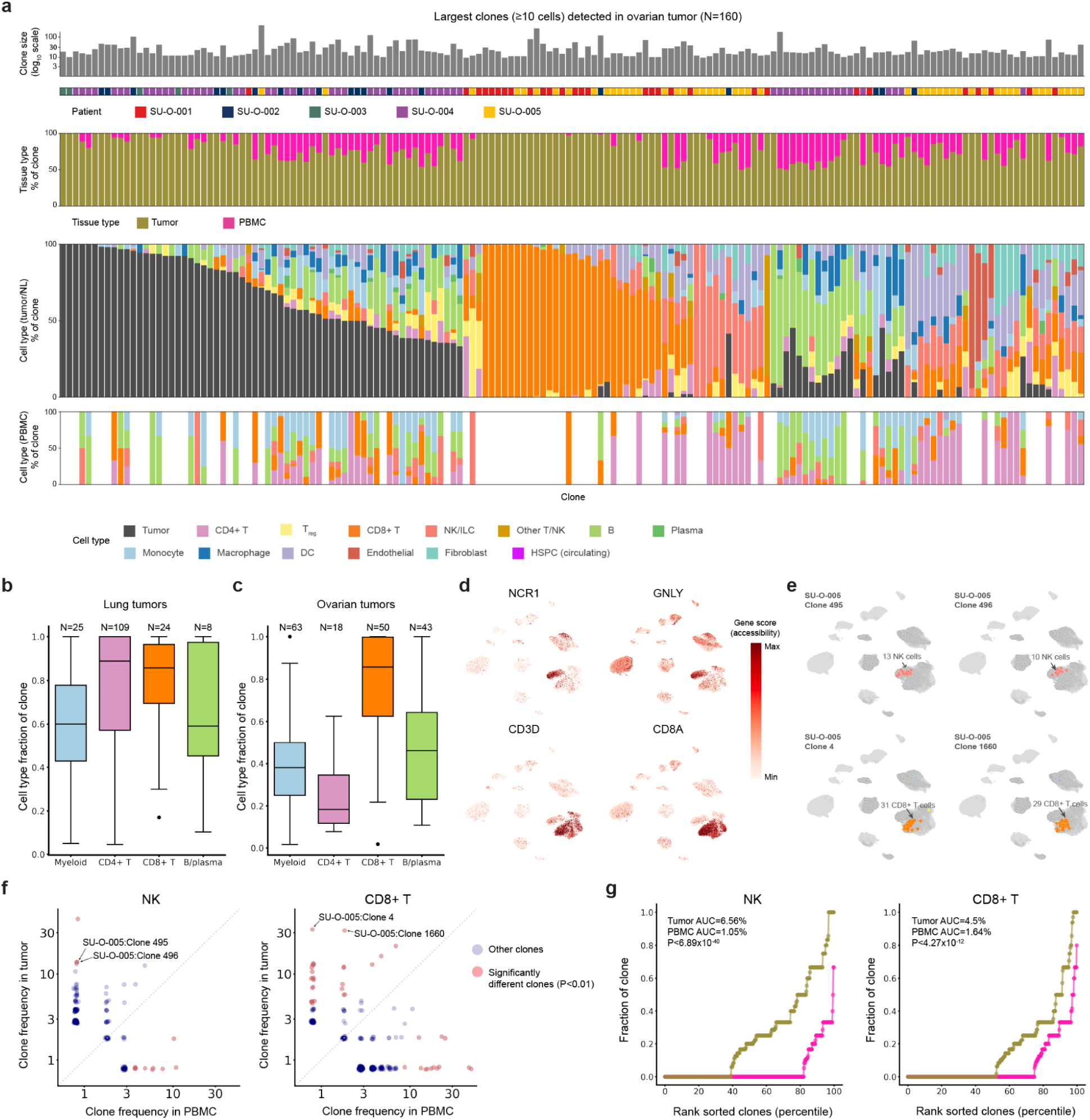
Additional clonal landscape analyses. (A) All clones with at least 10 cells detected in ovarian tumor samples. Each column represents a unique clone. (B and C) Clones with ≥5 cells of the indicated cell type are considered, and distribution of the indicated cell type’s clone fraction is plotted. For myeloid cells, monocytes, macrophages, and DCs are grouped. For CD4^+^ T cells, T_reg_s and other CD4^+^ T cells are grouped. N represents the number of clones considered for each indicated cell type. Analysis of lung tumor clones are displayed in (**B**) and ovarian tumor clones in (**C**). (D) Expression of NK and CD8^+^ T markers distinguish them on UMAP. (E) Representative clones capturing clonal expansion events of NK cells (top row) and CD8^+^ T cells (bottom row) in SU-O-005. For each clone, cells from the clone’s donor are highlighted with shaded circles, and cells assigned to that clone are colored by their cell type. (F) Scatterplots comparing clone frequencies of circulating cells with those infiltrating the tumor. The largest clones are locally expanded and minimally detected in the periphery. Significance is determined by Benjamini-Hochberg adjusted Fisher’s exact test. (G) Cumulative fraction of clone sizes for the indicated lymphoid cell types, split by tissue site. AUC corresponds to the overall clone size for the indicated cell type and tissue site. Kruskal-Wallis test.

**Supplemental Figure 5:**
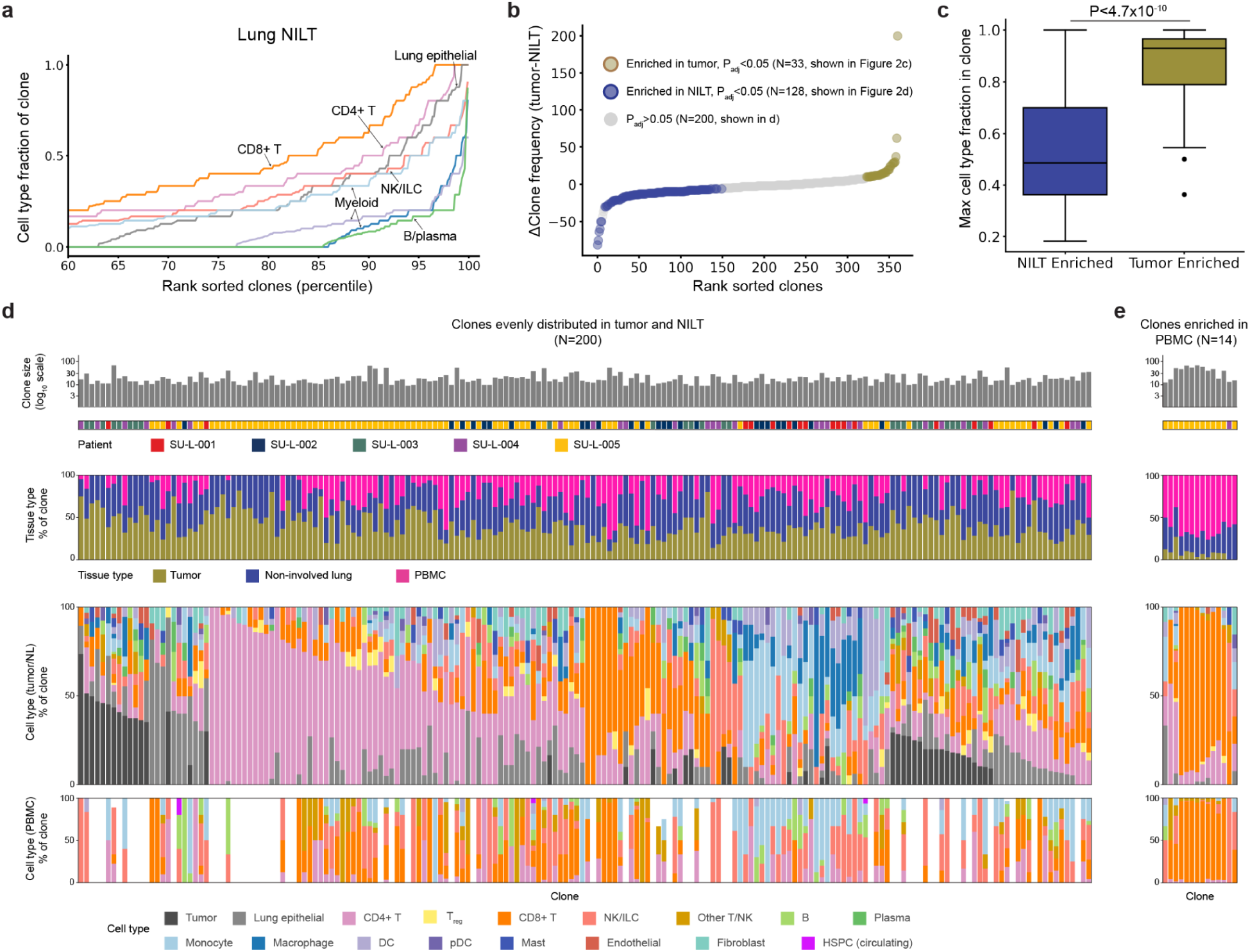
Comparative clonal analysis across tissue sites in NSCLC. (A) Cumulative fractions of clones stratified by cell type for cells from NILT samples. Clones with ≥5 cells are considered for this analysis. (B) Enrichment of clones in NILT or tumor. Significance is determined by Benjamini-Hochberg adjusted Fisher’s exact test against overall tissue site distribution for clones with at least 10 cells in tumor and NILT. (C) Comparison of dominant cell type fraction distribution between clones enriched in NILT and in tumor. Kruskal-Wallis test. (D and E) Heatmaps showing the fraction of all cell pairs belonging to the same clone and consisting of two cell types within ovarian tumor (**E**) and PBMC (**F**). Pairs were restricted to cells from the same donor.

**Supplemental Figure 6:**
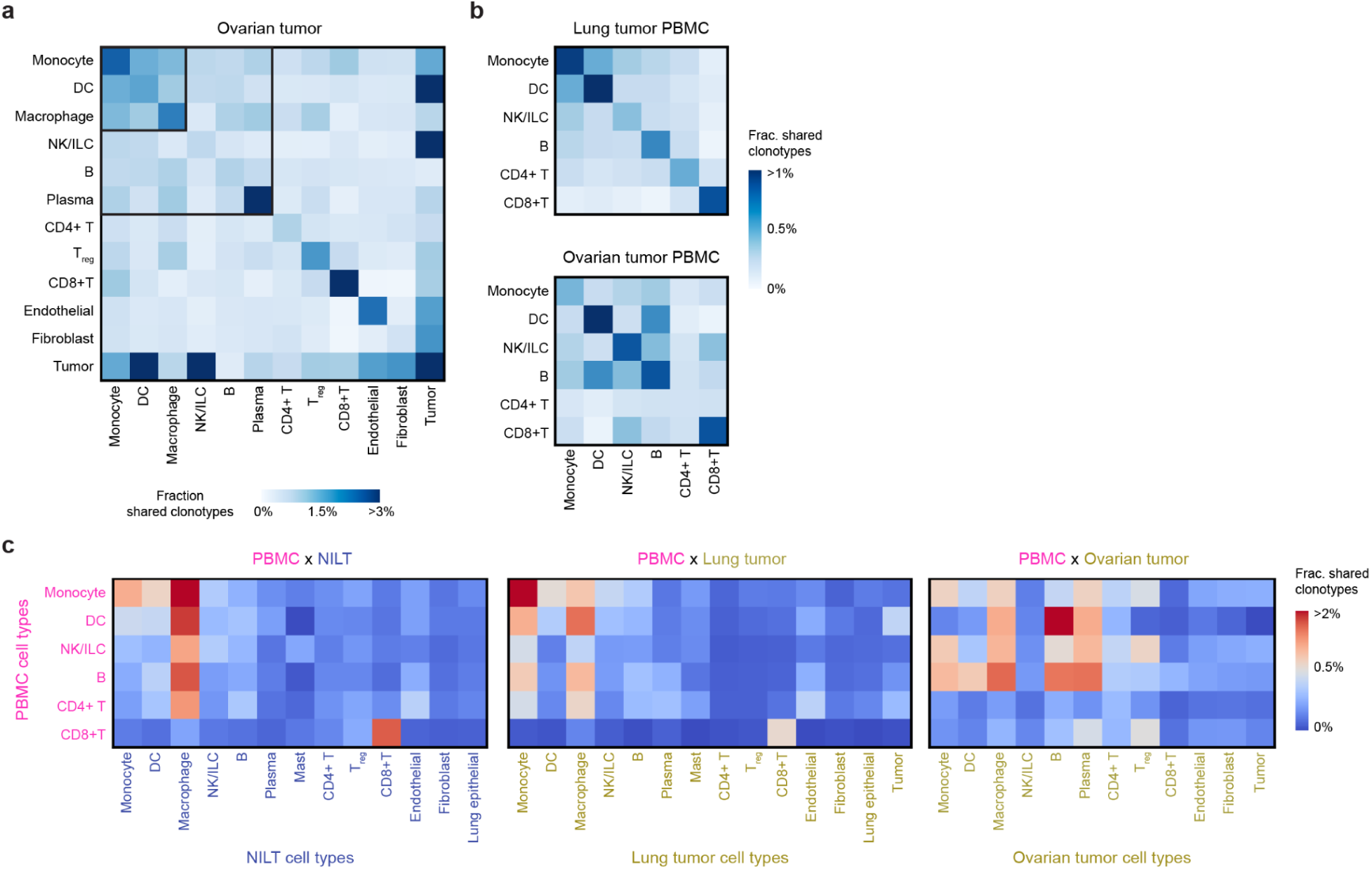
Additional cross-tissue clonal analysis. (A and B) Heatmaps showing the fraction of all cell pairs belonging to the same clone and consisting of two cell types within ovarian tumor (**A**) and PBMC (**B**). Pairs were restricted to cells from the same donor. (C) Heatmap showing the fraction of all cell pairs belonging to the same clone and consisting of a PBMC cell type and a solid tissue cell type.

**Supplemental Figure 7:**
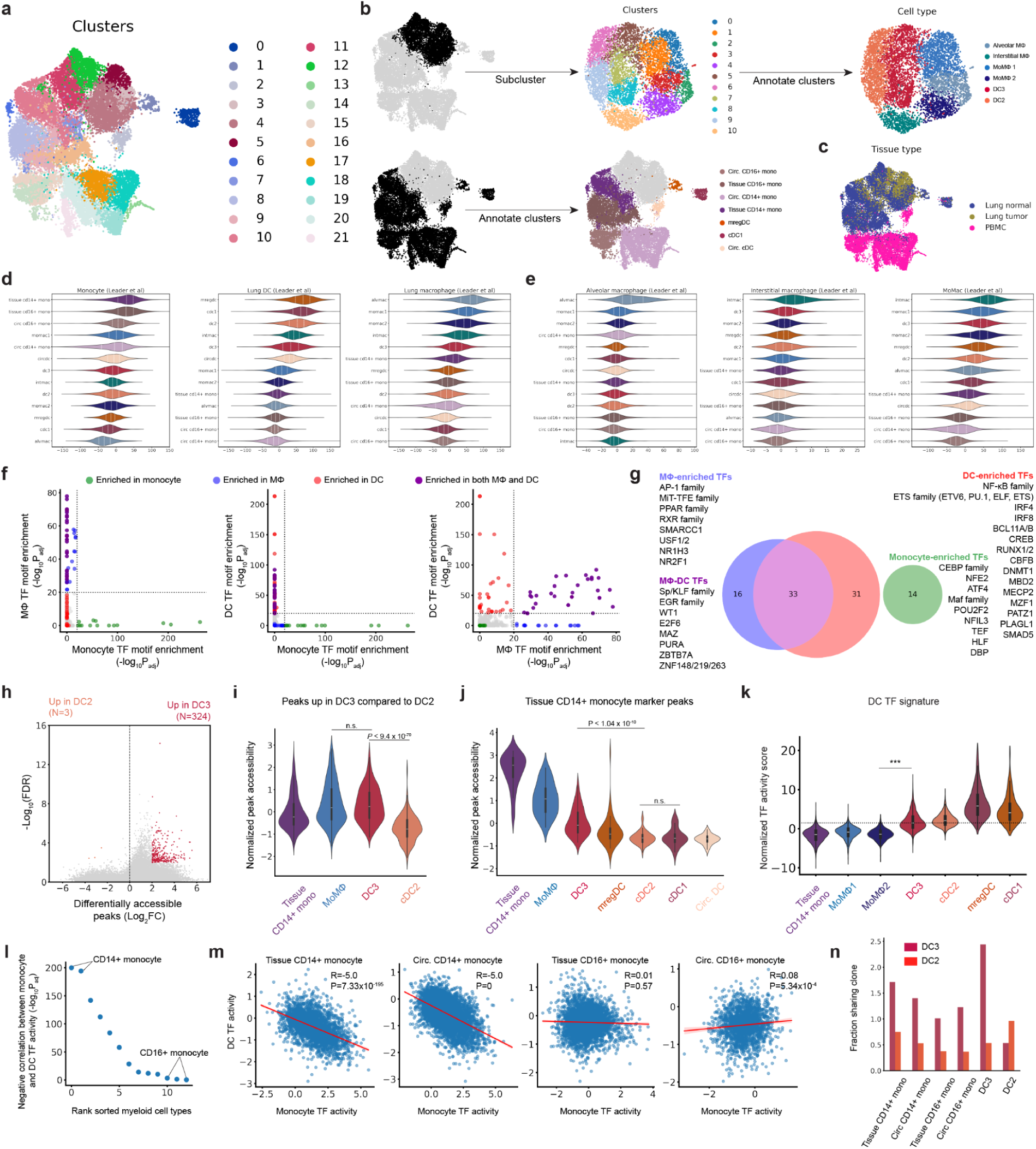
NSCLC myeloid annotation and epigenetic analysis. (A) UMAP of myeloid cells from PBMC, lung tumor, and NILT samples of patients with NSCLC, colored by their original cluster assignments. (B) Myeloid annotation scheme. Monocyte clusters separated clearly into CD14+ and CD16+ subsets, which were annotated without further subclustering. Macrophages and DCs were subclustered to achieve higher granularity and annotated based on subclustering results. (C) UMAP of cells colored by tissue sites. (D and E) Gene scores of published signatures derived from single-cell RNA and proteomic data. (F) TF motifs enriched in the marker peaks of three major MNP cell types. P-values are calculated from the Benjamini-Hochberg adjusted Wilcoxon signed-rank test. (G) Summary of TFs whose motifs are enriched in broad MNP cell types. A -log_10_P_adj_>20 cutoff was used. (H) Differentially accessible genomic regions in DC3 vs DC2. (I) Normalized sum accessibility of genomic regions significantly more accessible in DC3 compared to DC2 in indicated cell types. (J) Normalized sum accessibility of genomic regions significantly more accessible in CD14+ monocytes compared to other myeloid subtypes. (K) Average chromVAR motif deviation scores for DC TFs highlighted in Figure 3C. Kruskal-Wallis test. (L) Statistical significance of monocyte-DC TF motif accessibility correlation in all myeloid subtypes. CD14+ monocytes in tissue and circulation display the most significant negative correlation between monocyte and DC TF motif accessibilities. (M) Monocyte and DC TF activities are negatively correlated only in CD14+ monocytes but not in CD16+ monocytes. (N) Monocytes share more clonotypes with DC3 than with DC2.

**Supplemental Figure 8:**
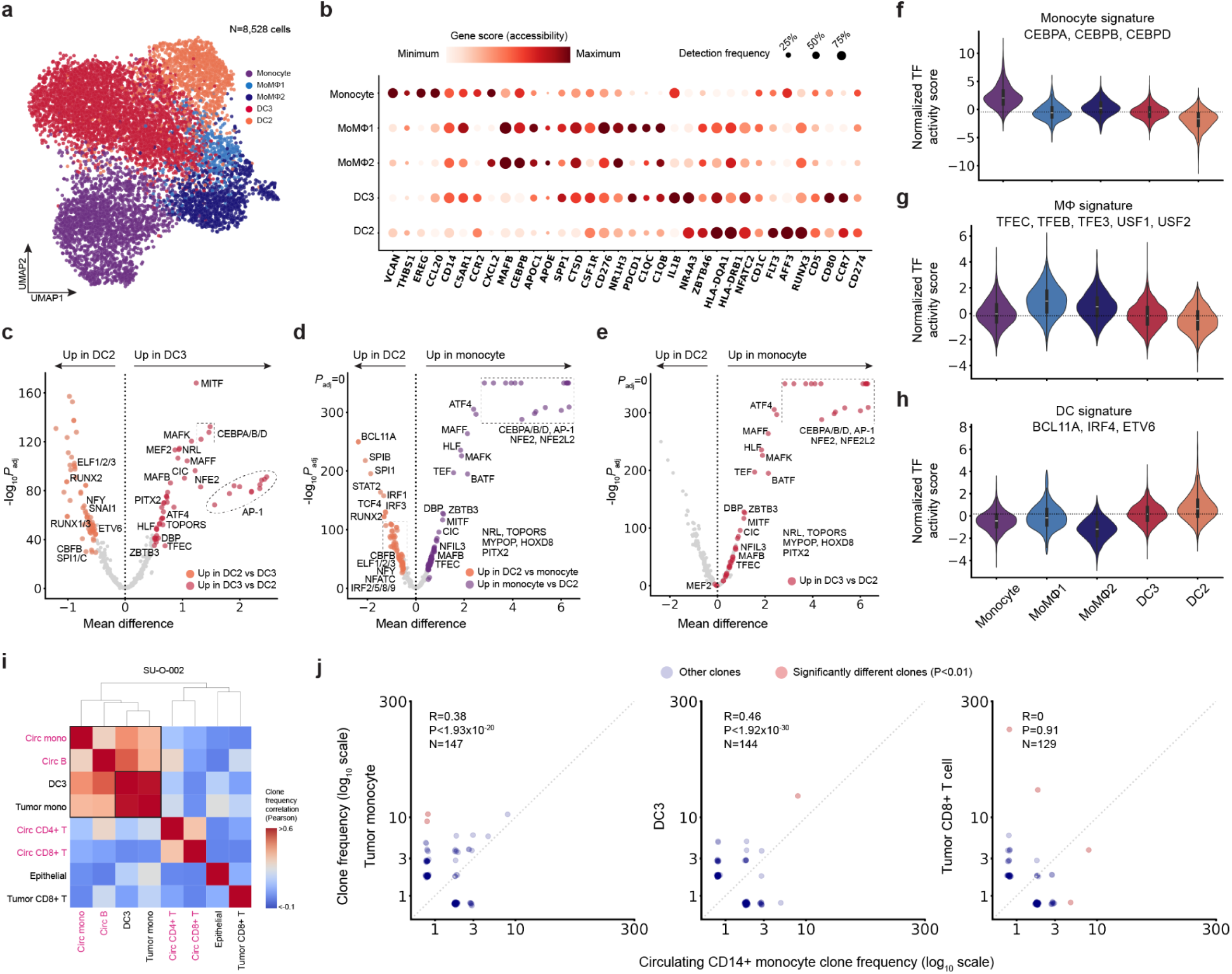
Ovarian myeloid epigenetic analysis. (A) UMAP of myeloid cells from ovarian tumor samples. (B) Column-scaled gene accessibility scores and detection frequencies for the indicated genes. (C, D, and E) Differentially active TF motifs between DC2 and DC3 (**C**) and DC2 and CD14+ monocytes (**D** and **E**). DC3-up TF motifs from (**D**) are highlighted in red in (**E**). P-values are calculated using the Benjamini-Hochberg adjusted Kruskal-Wallis test. (F, G, and H) Average chromVAR motif deviation scores for the indicated monocyte-, macrophage-, and DC-associated TFs. (I) Cell type-cell type clone frequency correlation across clones (≥5 cells across all samples). Color denotes correlation value, computed using Pearson’s ρ. Text labels of circulating PBMC cell types are colored pink. (J) Scatterplots comparing clone frequencies of circulating CD14+ monocyte and tissue CD14+ monocyte (left), DC3 (middle), and tumor CD8^+^ T cell (right). Significantly different clones with P<0.05 adjusted Fisher’s exact test are highlighted red.

**Supplemental Figure 9:**
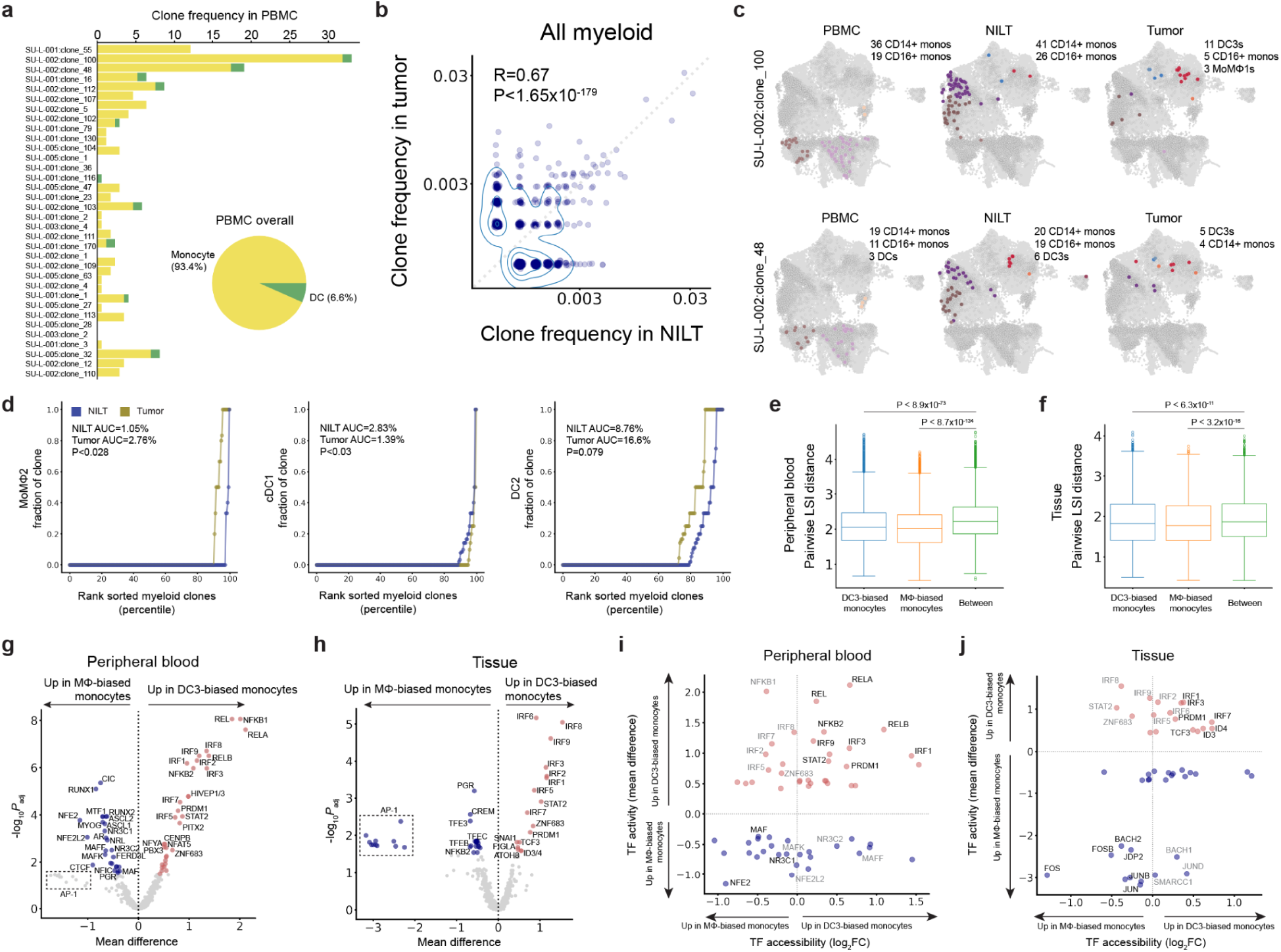
Epigenetic comparison of DC- and macrophage-biased myeloid clones. (A) Monocyte and DC proportions of largest myeloid clones (≥10 cells) in PBMC samples of patients with NSCLC. (B) Scatterplots comparing clone frequencies of myeloid cells in tumor with those in NILT. Contours visualize density. (C) Representative clones capturing myeloid cell type distribution across tissue sites. For each clone, cells from the clone’s donor are highlighted with shaded circles, and cells assigned to that clone are colored by their cell type. (D) Cumulative fraction of clone sizes for the indicated myeloid cell types, split by tissue site. AUC corresponds to the overall clone size for the indicated cell type and tissue site. Kruskal-Wallis test. (E and F) Epigenetic similarity as measured by distance in the LSI space within and between DC3-biased and macrophage-biased monocytes in circulation (**E**) and in tissue (**F**). Kruskal-Wallis test. (G and H) Differentially active TF motifs between DC3-biased and macrophage-biased monocytes in peripheral blood (**G**) and in tissue (**H**). P-values are calculated using the Benjamini-Hochberg adjusted Kruskal-Wallis test. (I and J) Joint comparison of TF gene body accessibility and inferred genome-wide TF activity to nominate TFs driving differential fate outcomes of monocytes in circulation (**I**) and in tissue (**J**). Select TFs that are both more accessible and motif accessibility are indicated in black, while TFs that only have increased motif accessibility are indicated in grey.

## Acknowledgements

We acknowledge the patients who volunteered to participate in these studies. We thank the members of the Satpathy Lab, L. Lanier, and Y. Lavin for stimulating discussions. A.T.S. was supported by a Career Award for Medical Scientists from the Burroughs Wellcome Fund, a Lloyd J. Old STAR Award from the Cancer Research Institute, and the Parker Institute for Cancer Immunotherapy. This work was supported by NIH grants P30CA008748 (C.A.L), R00HG012579 (C.A.L.), U01AT012984 (C.A.L.), and UM1HG012076 (A.T.S.; L.S.L.). L.S.L. is supported by the German Research Foundation (DFG) and an Emmy Noether fellowship (LU 2336/2-1). C.A.L. and A.T.S. are supported by the Michelson Medical Research Foundation. This work was supported by the Department of Pathology at Stanford University.

## Declaration of interests

V.L. is a consultant to Genesis Therapeutics and Cartography Biosciences. C.A.L is a consultant to Cartography Biosciences. A.T.S. is a founder of Immunai, Cartography Biosciences, Santa Ana Bio, and Prox Biosciences, an advisor to Zafrens and Wing Venture Capital, and receives research funding from Astellas.

## References

1. Pai, J.A., and Satpathy, A.T. (2021). High-throughput and single-cell T cell receptor sequencing technologies. Nat. Methods 18, 881–892.

2. Bradley, P., and Thomas, P.G. (2019). Using T cell receptor repertoires to understand the principles of adaptive immune recognition. Annu. Rev. Immunol. 37, 547–570.

3. Yost, K.E., Satpathy, A.T., Wells, D.K., Qi, Y., Wang, C., Kageyama, R., McNamara, K.L., Granja, J.M., Sarin, K.Y., Brown, R.A., et al. (2019). Clonal replacement of tumor-specific T cells following PD-1 blockade. Nat. Med. 25, 1251–1259.

4. Wu, T.D., Madireddi, S., de Almeida, P.E., Banchereau, R., Chen, Y.-J.J., Chitre, A.S., Chiang, E.Y., Iftikhar, H., O’Gorman, W.E., Au-Yeung, A., et al. (2020). Peripheral T cell expansion predicts tumour infiltration and clinical response. Nature 579, 274–278.

5. Pai, J.A., Hellmann, M.D., Sauter, J.L., Mattar, M., Rizvi, H., Woo, H.J., Shah, N., Nguyen, E.M., Uddin, F.Z., Quintanal-Villalonga, A., et al. (2023). Lineage tracing reveals clonal progenitors and long-term persistence of tumor-specific T cells during immune checkpoint blockade. Cancer Cell 41, 776–790.e7.

6. Cassetta, L., and Pollard, J.W. (2018). Targeting macrophages: therapeutic approaches in cancer. Nat. Rev. Drug Discov. 17, 887–904.

7. van Vlerken-Ysla, L., Tyurina, Y.Y., Kagan, V.E., and Gabrilovich, D.I. (2023). Functional states of myeloid cells in cancer. Cancer Cell 41, 490–504.

8. Lavin, Y., Kobayashi, S., Leader, A., Amir, E.-A.D., Elefant, N., Bigenwald, C., Remark, R., Sweeney, R., Becker, C.D., Levine, J.H., et al. (2017). Innate Immune Landscape in Early Lung Adenocarcinoma by Paired Single-Cell Analyses. Cell 169, 750–765.e17.

9. Zhang, Q., He, Y., Luo, N., Patel, S.J., Han, Y., Gao, R., Modak, M., Carotta, S., Haslinger, C., Kind, D., et al. (2019). Landscape and dynamics of single immune cells in hepatocellular carcinoma. Cell 179, 829–845.e20.

10. Steele, N.G., Carpenter, E.S., Kemp, S.B., Sirihorachai, V.R., The, S., Delrosario, L., Lazarus, J., Amir, E.-A.D., Gunchick, V., Espinoza, C., et al. (2020). Multimodal mapping of the tumor and peripheral blood immune landscape in human pancreatic cancer. Nat. Cancer 1, 1097–1112.

11. Krishna, C., DiNatale, R.G., Kuo, F., Srivastava, R.M., Vuong, L., Chowell, D., Gupta, S., Vanderbilt, C., Purohit, T.A., Liu, M., et al. (2021). Single-cell sequencing links multiregional immune landscapes and tissue-resident T cells in ccRCC to tumor topology and therapy efficacy. Cancer Cell 39, 662–677.e6.

12. Casanova-Acebes, M., Dalla, E., Leader, A.M., LeBerichel, J., Nikolic, J., Morales, B.M., Brown, M., Chang, C., Troncoso, L., Chen, S.T., et al. (2021). Tissue-resident macrophages provide a pro-tumorigenic niche to early NSCLC cells. Nature 595, 578–584.

13. Nalio Ramos, R., Missolo-Koussou, Y., Gerber-Ferder, Y., Bromley, C.P., Bugatti, M., Núñez, N.G., Tosello Boari, J., Richer, W., Menger, L., Denizeau, J., et al. (2022). Tissue-resident FOLR2+ macrophages associate with CD8+ T cell infiltration in human breast cancer. Cell 185, 1189–1207.e25.

14. Larsson, N.-G. (2010). Somatic mitochondrial DNA mutations in mammalian aging. Annu. Rev. Biochem. 79, 683–706.

15. Stewart, J.B., and Chinnery, P.F. (2015). The dynamics of mitochondrial DNA heteroplasmy: implications for human health and disease. Nat. Rev. Genet. 16, 530–542.

16. Ludwig, L.S., Lareau, C.A., Ulirsch, J.C., Christian, E., Muus, C., Li, L.H., Pelka, K., Ge, W., Oren, Y., Brack, A., et al. (2019). Lineage Tracing in Humans Enabled by Mitochondrial Mutations and Single-Cell Genomics. Cell 176, 1325–1339.e22.

17. Lareau, C.A., Ludwig, L.S., Muus, C., Gohil, S.H., Zhao, T., Chiang, Z., Pelka, K., Verboon, J.M., Luo, W., Christian, E., et al. (2021). Massively parallel single-cell mitochondrial DNA genotyping and chromatin profiling. Nat. Biotechnol. 39, 451–461.

18. Weng, C., Yu, F., Yang, D., Poeschla, M., Liggett, L.A., Jones, M.G., Qiu, X., Wahlster, L., Caulier, A., Hussmann, J.A., et al. (2024). Deciphering cell states and genealogies of human hematopoiesis.

19. Lareau, C.A., Liu, V., Muus, C., Praktiknjo, S.D., Nitsch, L., Kautz, P., Sandor, K., Yin, Y., Gutierrez, J.C., Pelka, K., et al. (2023). Mitochondrial single-cell ATAC-seq for high-throughput multi-omic detection of mitochondrial genotypes and chromatin accessibility. Nat. Protoc. 18, 1416–1440.

20. Rückert, T., Lareau, C.A., Mashreghi, M.-F., Ludwig, L.S., and Romagnani, C. (2022). Clonal expansion and epigenetic inheritance of long-lasting NK cell memory. Nat. Immunol. 23, 1551–1563.

21. An, J., Nam, C.H., Kim, R., Lee, Y., Won, H., Park, S., Lee, W.H., Park, H., Yoon, C.J., An, Y., et al. (2024). Mitochondrial DNA mosaicism in normal human somatic cells. Nat. Genet. 56, 1665–1677.

22. Jenuth, J.P., Peterson, A.C., Fu, K., and Shoubridge, E.A. (1996). Random genetic drift in the female germline explains the rapid segregation of mammalian mitochondrial DNA. Nat. Genet. 14, 146–151.

23. De Rop, F.V., Hulselmans, G., Flerin, C., Soler-Vila, P., Rafels, A., Christiaens, V., González-Blas, C.B., Marchese, D., Caratù, G., Poovathingal, S., et al. (2024). Systematic benchmarking of single-cell ATAC-sequencing protocols. Nat. Biotechnol. 42, 916–926.

24. Borcherding, N., and Brestoff, J.R. (2023). The power and potential of mitochondria transfer. Nature 623, 283–291.

25. Miller, T.E., Lareau, C.A., Verga, J.A., DePasquale, E.A.K., Liu, V., Ssozi, D., Sandor, K., Yin, Y., Ludwig, L.S., El Farran, C.A., et al. (2022). Mitochondrial variant enrichment from high-throughput single-cell RNA sequencing resolves clonal populations. Nat. Biotechnol. 40, 1030–1034.

26. Wu, F., Fan, J., He, Y., Xiong, A., Yu, J., Li, Y., Zhang, Y., Zhao, W., Zhou, F., Li, W., et al. (2021). Single-cell profiling of tumor heterogeneity and the microenvironment in advanced non-small cell lung cancer. Nat. Commun. 12, 2540.

27. Leader, A.M., Grout, J.A., Maier, B.B., Nabet, B.Y., Park, M.D., Tabachnikova, A., Chang, C., Walker, L., Lansky, A., Le Berichel, J., et al. (2021). Single-cell analysis of human non-small cell lung cancer lesions refines tumor classification and patient stratification. Cancer Cell 39, 1594–1609.e12.

28. Jardim, D.L., Goodman, A., de Melo Gagliato, D., and Kurzrock, R. (2021). The challenges of tumor mutational burden as an immunotherapy biomarker. Cancer Cell 39, 154–173.

29. Kahn, B.M., Lucas, A., Alur, R.G., Wengyn, M.D., Schwartz, G.W., Li, J., Sun, K., Maurer, H.C., Olive, K.P., Faryabi, R.B., et al. (2021). The vascular landscape of human cancer. J. Clin. Invest. 131. 10.1172/JCI136655.

30. Ghoneum, A., Afify, H., Salih, Z., Kelly, M., and Said, N. (2018). Role of tumor microenvironment in ovarian cancer pathobiology. Oncotarget 9, 22832–22849.

31. Hornburg, M., Desbois, M., Lu, S., Guan, Y., Lo, A.A., Kaufman, S., Elrod, A., Lotstein, A., DesRochers, T.M., Munoz-Rodriguez, J.L., et al. (2021). Single-cell dissection of cellular components and interactions shaping the tumor immune phenotypes in ovarian cancer. Cancer Cell 39, 928–944.e6.

32. Regner, M.J., Wisniewska, K., Garcia-Recio, S., Thennavan, A., Mendez-Giraldez, R., Malladi, V.S., Hawkins, G., Parker, J.S., Perou, C.M., Bae-Jump, V.L., et al. (2021). A multi-omic single-cell landscape of human gynecologic malignancies. Mol. Cell 81, 4924–4941.e10.

33. Sun, J.C., Beilke, J.N., and Lanier, L.L. (2009). Adaptive immune features of natural killer cells. Nature 457, 557–561.

34. Hammer, Q., Rückert, T., Borst, E.M., Dunst, J., Haubner, A., Durek, P., Heinrich, F., Gasparoni, G., Babic, M., Tomic, A., et al. (2018). Peptide-specific recognition of human cytomegalovirus strains controls adaptive natural killer cells. Nat. Immunol. 19, 453–463.

35. Béziat, V., Liu, L.L., Malmberg, J.-A., Ivarsson, M.A., Sohlberg, E., Björklund, A.T., Retière, C., Sverremark-Ekström, E., Traherne, J., Ljungman, P., et al. (2013). NK cell responses to cytomegalovirus infection lead to stable imprints in the human KIR repertoire and involve activating KIRs. Blood 121, 2678–2688.

36. Strauss-Albee, D.M., Fukuyama, J., Liang, E.C., Yao, Y., Jarrell, J.A., Drake, A.L., Kinuthia, J., Montgomery, R.R., John-Stewart, G., Holmes, S., et al. (2015). Human NK cell repertoire diversity reflects immune experience and correlates with viral susceptibility. Sci. Transl. Med. 7. 10.1126/scitranslmed.aac5722.

37. MacLean, A.J., Bonifacio, J.P.P.L., Oram, S.L., Mohsen, M.O., Bachmann, M.F., and Arnon, T.I. (2024). Regulation of pulmonary plasma cell responses during secondary infection with influenza virus. J. Exp. Med. 221. 10.1084/jem.20232014.

38. Hei Yu, K.K., Abou-Mrad, Z., Törkenczy, K., Schulze, I., Gantchev, J., Baquer, G., Hopland, K., Bander, E.D., Tosi, U., Brennan, C., et al. (2025). A pathogenic subpopulation of human glioma associated macrophages linked to glioma progression. bioRxivorg. 10.1101/2025.02.12.637857.

39. Park, M.D., Reyes-Torres, I., LeBerichel, J., Hamon, P., LaMarche, N.M., Hegde, S., Belabed, M., Troncoso, L., Grout, J.A., Magen, A., et al. (2023). TREM2 macrophages drive NK cell paucity and dysfunction in lung cancer. Nat. Immunol. 24, 792–801.

40. Villani, A.-C., Satija, R., Reynolds, G., Sarkizova, S., Shekhar, K., Fletcher, J., Griesbeck, M., Butler, A., Zheng, S., Lazo, S., et al. (2017). Single-cell RNA-seq reveals new types of human blood dendritic cells, monocytes, and progenitors. Science 356. 10.1126/science.aah4573.

41. Bourdely, P., Anselmi, G., Vaivode, K., Ramos, R., Missolo-Koussou, Y., Hidalgo, S., Tosselo, J., Núñez, N., Richer, W., Vincent-Salomon, A., et al. (2020). Transcriptional and Functional Analysis of CD1c+ Human Dendritic Cells Identifies a CD163+ Subset Priming CD8+CD103+ T Cells. Immunity 53, 335–352.e8.

42. Brown, C.C., Gudjonson, H., Pritykin, Y., Deep, D., Lavallée, V.-P., Mendoza, A., Fromme, R., Mazutis, L., Ariyan, C., Leslie, C., et al. (2019). Transcriptional basis of mouse and human dendritic cell heterogeneity. Cell 179, 846–863.e24.

43. Dutertre, C.-A., Becht, E., Irac, S.E., Khalilnezhad, A., Narang, V., Khalilnezhad, S., Ng, P.Y., van den Hoogen, L.L., Leong, J.Y., Lee, B., et al. (2019). Single-cell analysis of human mononuclear phagocytes reveals subset-defining markers and identifies circulating inflammatory dendritic cells. Immunity 51, 573–589.e8.

44. Maier, B., Leader, A.M., Chen, S.T., Tung, N., Chang, C., LeBerichel, J., Chudnovskiy, A., Maskey, S., Walker, L., Finnigan, J.P., et al. (2020). A conserved dendritic-cell regulatory program limits antitumour immunity. Nature 580, 257–262.

45. Zilionis, R., Engblom, C., Pfirschke, C., Savova, V., Zemmour, D., Saatcioglu, H.D., Krishnan, I., Maroni, G., Meyerovitz, C.V., Kerwin, C.M., et al. (2019). Single-cell transcriptomics of human and mouse lung cancers reveals conserved myeloid populations across individuals and species. Immunity 50, 1317–1334.e10.

46. Granja, J.M., Corces, M.R., Pierce, S.E., Bagdatli, S.T., Choudhry, H., Chang, H.Y., and Greenleaf, W.J. (2021). ArchR is a scalable software package for integrative single-cell chromatin accessibility analysis. Nat. Genet. 53, 403–411.

47. Zhang, D.E., Zhang, P., Wang, N.D., Hetherington, C.J., Darlington, G.J., and Tenen, D.G. (1997). Absence of granulocyte colony-stimulating factor signaling and neutrophil development in CCAAT enhancer binding protein alpha-deficient mice. Proc. Natl. Acad. Sci. U. S. A. 94, 569–574.

48. Kim, S., Chen, J., Ou, F., Liu, T.-T., Jo, S., Gillanders, W.E., Murphy, T.L., and Murphy, K.M. (2024). Transcription factor C/EBPα is required for the development of Ly6C hi monocytes but not Ly6C lo monocytes. Proc. Natl. Acad. Sci. U.S.A. 121. 10.1073/pnas.2315659121.

49. Kirschenbaum, D., Xie, K., Ingelfinger, F., Katzenelenbogen, Y., Abadie, K., Look, T., Sheban, F., Phan, T.S., Li, B., Zwicky, P., et al. (2024). Time-resolved single-cell transcriptomics defines immune trajectories in glioblastoma. Cell 187, 149–165.e23.

50. Dekkers, K.F., Neele, A.E., Jukema, J.W., Heijmans, B.T., and de Winther, M.P.J. (2019). Human monocyte-to-macrophage differentiation involves highly localized gain and loss of DNA methylation at transcription factor binding sites. Epigenetics Chromatin 12, 34.

51. Phanstiel, D.H., Van Bortle, K., Spacek, D., Hess, G.T., Shamim, M.S., Machol, I., Love, M.I., Aiden, E.L., Bassik, M.C., and Snyder, M.P. (2017). Static and Dynamic DNA Loops form AP-1-Bound Activation Hubs during Macrophage Development. Mol. Cell 67, 1037–1048.e6.

52. Devalaraja, S., To, T.K.J., Folkert, I.W., Natesan, R., Alam, M.Z., Li, M., Tada, Y., Budagyan, K., Dang, M.T., Zhai, L., et al. (2020). Tumor-derived retinoic acid regulates intratumoral monocyte differentiation to promote immune suppression. Cell 180, 1098–1114.e16.

53. Sharma, S.M., Bronisz, A., Hu, R., Patel, K., Mansky, K.C., Sif, S., and Ostrowski, M.C. (2007). MITF and PU.1 recruit p38 MAPK and NFATc1 to target genes during osteoclast differentiation. J. Biol. Chem. 282, 15921–15929.

54. Liao, X., Sharma, N., Kapadia, F., Zhou, G., Lu, Y., Hong, H., Paruchuri, K., Mahabeleshwar, G.H., Dalmas, E., Venteclef, N., et al. (2011). Krüppel-like factor 4 regulates macrophage polarization. J. Clin. Invest. 121, 2736–2749.

55. A-González, N., and Castrillo, A. (2011). Liver X receptors as regulators of macrophage inflammatory and metabolic pathways. Biochim. Biophys. Acta 1812, 982–994.

56. Goudot, C., Coillard, A., Villani, A.-C., Gueguen, P., Cros, A., Sarkizova, S., Tang-Huau, T.-L., Bohec, M., Baulande, S., Hacohen, N., et al. (2017). Aryl hydrocarbon receptor controls monocyte differentiation into dendritic cells versus macrophages. Immunity 47, 582–596.e6.

57. Trizzino, M., Zucco, A., Deliard, S., Wang, F., Barbieri, E., Veglia, F., Gabrilovich, D., and Gardini, A. (2021). EGR1 is a gatekeeper of inflammatory enhancers in human macrophages. Sci. Adv. 7. 10.1126/sciadv.aaz8836.

58. Moore, K.J., Rosen, E.D., Fitzgerald, M.L., Randow, F., Andersson, L.P., Altshuler, D., Milstone, D.S., Mortensen, R.M., Spiegelman, B.M., and Freeman, M.W. (2001). The role of PPAR-gamma in macrophage differentiation and cholesterol uptake. Nat. Med. 7, 41–47.

59. Lavin, Y., Winter, D., Blecher-Gonen, R., David, E., Keren-Shaul, H., Merad, M., Jung, S., and Amit, I. (2014). Tissue-resident macrophage enhancer landscapes are shaped by the local microenvironment. Cell 159, 1312–1326.

60. Murphy, T.L., Grajales-Reyes, G.E., Wu, X., Tussiwand, R., Briseño, C.G., Iwata, A., Kretzer, N.M., Durai, V., and Murphy, K.M. (2016). Transcriptional control of dendritic cell development. Annu. Rev. Immunol. 34, 93–119.

61. Satpathy, A.T., Briseño, C.G., Cai, X., Michael, D.G., Chou, C., Hsiung, S., Bhattacharya, D., Speck, N.A., and Egawa, T. (2014). Runx1 and Cbfβ regulate the development of Flt3+ dendritic cell progenitors and restrict myeloproliferative disorder. Blood 123, 2968–2977.

62. Villar, J., Cros, A., De Juan, A., Alaoui, L., Bonte, P.-E., Lau, C.M., Tiniakou, I., Reizis, B., and Segura, E. (2023). ETV3 and ETV6 enable monocyte differentiation into dendritic cells by repressing macrophage fate commitment. Nat. Immunol. 24, 84–95.

63. Shih, V.F.-S., Davis-Turak, J., Macal, M., Huang, J.Q., Ponomarenko, J., Kearns, J.D., Yu, T., Fagerlund, R., Asagiri, M., Zuniga, E.I., et al. (2012). Control of RelB during dendritic cell activation integrates canonical and noncanonical NF-κB pathways. Nat. Immunol. 13, 1162–1170.

64. Alder, J.K., Georgantas, R.W., 3rd, Hildreth, R.L., Kaplan, I.M., Morisot, S., Yu, X., McDevitt, M., and Civin, C.I. (2008). Kruppel-like factor 4 is essential for inflammatory monocyte differentiation in vivo. J. Immunol. 180, 5645–5652.

65. Feinberg, M.W., Wara, A.K., Cao, Z., Lebedeva, M.A., Rosenbauer, F., Iwasaki, H., Hirai, H., Katz, J.P., Haspel, R.L., Gray, S., et al. (2007). The Kruppel-like factor KLF4 is a critical regulator of monocyte differentiation. EMBO J. 26, 4138–4148.

66. Tussiwand, R., Everts, B., Grajales-Reyes, G.E., Kretzer, N.M., Iwata, A., Bagaitkar, J., Wu, X., Wong, R., Anderson, D.A., Murphy, T.L., et al. (2015). Klf4 expression in conventional dendritic cells is required for T helper 2 cell responses. Immunity 42, 916–928.

67. Carter, J.H., and Tourtellotte, W.G. (2007). Early growth response transcriptional regulators are dispensable for macrophage differentiation. J. Immunol. 178, 3038–3047.

68. Guilliams, M., Mildner, A., and Yona, S. (2018). Developmental and functional heterogeneity of monocytes. Immunity 49, 595–613.

69. Jakubzick, C.V., Randolph, G.J., and Henson, P.M. (2017). Monocyte differentiation and antigen-presenting functions. Nat. Rev. Immunol. 17, 349–362.

70. De Zuani, M., Xue, H., Park, J.S., Dentro, S.C., Seferbekova, Z., Tessier, J., Curras-Alonso, S., Hadjipanayis, A., Athanasiadis, E.I., Gerstung, M., et al. (2024). Single-cell and spatial transcriptomics analysis of non-small cell lung cancer. Nat. Commun. 15, 4388.

71. Kusiak, A., and Brady, G. (2022). Bifurcation of signalling in human innate immune pathways to NF-kB and IRF family activation. Biochem. Pharmacol. 205, 115246.

72. Kim, S.J. (2015). Immunological function of Blimp-1 in dendritic cells and relevance to autoimmune diseases. Immunol. Res. 63, 113–120.

73. Deng, Z., Loyher, P.-L., Lazarov, T., Li, L., Shen, Z., Bhinder, B., Yang, H., Zhong, Y., Alberdi, A., Massague, J., et al. (2024). The nuclear factor ID3 endows macrophages with a potent anti-tumour activity. Nature 626, 864–873.

74. Kobayashi, E.H., Suzuki, T., Funayama, R., Nagashima, T., Hayashi, M., Sekine, H., Tanaka, N., Moriguchi, T., Motohashi, H., Nakayama, K., et al. (2016). Nrf2 suppresses macrophage inflammatory response by blocking proinflammatory cytokine transcription. Nat. Commun. 7, 11624.

75. Zezulin, A.U., Ye, D., Howell, E., Yen, D., Bresciani, E., Diemer, J., Ren, J.-G., Ahmad, M.H., Castilla, L.H., Touw, I.P., et al. (2023). RUNX1 is required in granulocyte-monocyte progenitors to attenuate inflammatory cytokine production by neutrophils. bioRxivorg. 10.1101/2023.01.27.525911.

76. Bellissimo, D.C., Chen, C.-H., Zhu, Q., Bagga, S., Lee, C.-T., He, B., Wertheim, G.B., Jordan, M., Tan, K., Worthen, G.S., et al. (2020). Runx1 negatively regulates inflammatory cytokine production by neutrophils in response to Toll-like receptor signaling. Blood Adv. 4, 1145–1158.

77. Tomimatsu, M., Matsumoto, K., Ashizuka, M., Kumagai, S., Tanaka, S., Nakae, T., Yokota, K., Kominami, S., Kajiura, R., Okuzaki, D., et al. (2022). Myeloid cell-specific ablation of Runx2 gene exacerbates post-infarct cardiac remodeling. Sci. Rep. 12, 16656.

78. Cao, S., Liu, J., Song, L., and Ma, X. (2005). The protooncogene c-Maf is an essential transcription factor for IL-10 gene expression in macrophages. J. Immunol. 174, 3484–3492.

79. Liu, M., Tong, Z., Ding, C., Luo, F., Wu, S., Wu, C., Albeituni, S., He, L., Hu, X., Tieri, D., et al. (2020). Transcription factor c-Maf is a checkpoint that programs macrophages in lung cancer. J. Clin. Invest. 130, 2081–2096.

80. Cao, S., Liu, J., Chesi, M., Bergsagel, P.L., Ho, I.-C., Donnelly, R.P., and Ma, X. (2002). Differential regulation of IL-12 and IL-10 gene expression in macrophages by the basic leucine zipper transcription factor c-Maf fibrosarcoma. J. Immunol. 169, 5715–5725.

81. Liu, Z., Wang, H., Li, Z., Dress, R.J., Zhu, Y., Zhang, S., De Feo, D., Kong, W.T., Cai, P., Shin, A., et al. (2023). Dendritic cell type 3 arises from Ly6C+ monocyte-dendritic cell progenitors. Immunity 56, 1761–1777.e6.

82. Hiam-Galvez, K.J., Allen, B.M., and Spitzer, M.H. (2021). Systemic immunity in cancer. Nat. Rev. Cancer 21, 345–359.

83. Park, S.H., Kang, K., Giannopoulou, E., Qiao, Y., Kang, K., Kim, G., Park-Min, K.-H., and Ivashkiv, L.B. (2017). Type I interferons and the cytokine TNF cooperatively reprogram the macrophage epigenome to promote inflammatory activation. Nat. Immunol. 18, 1104–1116.

84. Sheu, K.M., and Hoffmann, A. (2022). Functional hallmarks of healthy macrophage responses: Their regulatory basis and disease relevance. Annu. Rev. Immunol. 40, 295–321.

85. Hao, Y., Stuart, T., Kowalski, M.H., Choudhary, S., Hoffman, P., Hartman, A., Srivastava, A., Molla, G., Madad, S., Fernandez-Granda, C., et al. (2024). Dictionary learning for integrative, multimodal and scalable single-cell analysis. Nat. Biotechnol. 42, 293–304.

86. Schep, A.N., Wu, B., Buenrostro, J.D., and Greenleaf, W.J. (2017). chromVAR: inferring transcription-factor-associated accessibility from single-cell epigenomic data. Nat. Methods 14, 975–978.

